# Cell Cycle-Dependent Chromatin Motion: A Role for DNA Content Doubling Over Cohesion

**DOI:** 10.64898/2026.03.19.712877

**Authors:** Martin Rey-Millet, Léa Costes, Erwan Le-Floch, Habib Ayoub, Quentin Saccomani, Manoel Manghi, Kerstin Bystricky

## Abstract

The spatiotemporal organisation of chromatin in the eukaryotic nucleus is fundamental for genome regulation. Chromatin undergoes rapid remodelling and rearrangements within minutes, altering its diffusion properties. Considering the tight coupling between genome function and nuclear architecture, a key question is how chromatin dynamics adapt to or promote nuclear processes. To elucidate the underlying physical principles, we employed High-resolution Diffusion mapping (Hi-D) to track chromatin motion throughout interphase in live human cells.

Our analysis, that considers both diffusive motion and drift generated by active forces, revealed that chromatin dynamics are heterogeneous, with distinct behaviours in different subnuclear zones. Notably, both diffusive and processive contributions to chromatin motion progressively decrease from G1 to G2 phase, with this reduction occurring uniformly across all subzones. This suggests a global mechanism driving the observed decrease in chromatin mobility during cell cycle progression.

By combining genetic knockout experiments and polymer modelling, we demonstrate that the doubling of DNA content, rather than cohesin-mediated sister chromatid entrapment, is responsible for the gradual decrease in chromatin motion during the cell cycle in human nuclei. These findings provide new insights into the physical and functional organisation of chromatin and its regulation during cellular proliferation.

**GRAPHICAL ABSTRACT:** 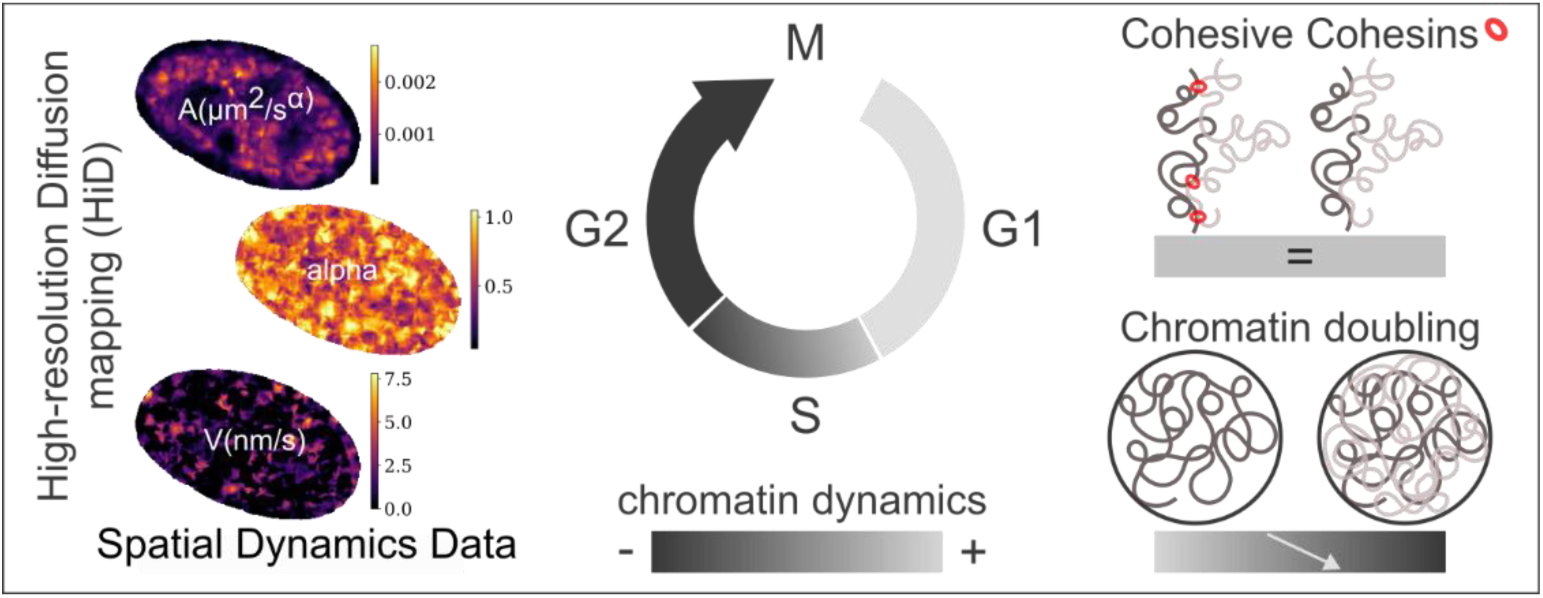

## INTRODUCTION

DNA in eukaryotes is organised at different scales, from the 10 nm chromatin fibre to chromosome territories. This three-dimensional (3D) genome architecture in interphase nuclei is essential for packing the long DNA molecule in the nucleus and allowing spatial variation in concentration of proteins and nucleic acids necessary to regulate transcription, DNA repair, replication, or ribosome biogenesis (1, 2). The development of genome wide chromosome conformation capture techniques and statistical modelling of measured average contact frequencies (3) has enabled a detailed exploration and understanding of 3D nuclear architecture. However, such techniques require cell fixation resulting snapshots of the spatial organisation at a given moment and therefore miss the parameter ‘time’ (4, 5). Yet, plasticity in the organisation of cellular processes is an essential biological feature to adapt and respond efficiently to changes.

Indeed, chromatin is highly mobile. Early experiments in yeast, tracking single chromatin foci over minutes, revealed the dynamic nature of labelled genome segments (6, 7), soon enriched by other observations in various eukaryotic systems and at different spatio-temporal scales (8–11). From a physical point of view, chromatin can be considered as a polymer, and tracking of local chromatin in living cells generally exhibits slow diffusion dynamics described by the anomalous sub-diffusion Rouse model (12–14). Nonetheless, free diffusion or active transport have also been observed (15).

There is a clear relationship between structure and function of chromatin at the scale of the entire nucleus (2). For instance, dense heterochromatin, such as lamina-associated domains (LAD) and nucleolus-associated domains (NAD), is preferentially located near the nuclear and nucleolar periphery, while euchromatin and sites of active transcription are mostly found in the nuclear interior. Activation of transcription has been shown to induce architectural rearrangements in chromatin structure (16, 17). Given the relation between genome function and architecture, one could ask how chromatin dynamics respond to or facilitate nuclear processes and what the contribution of active forces is. One line of evidence that emerged early in the field is the interdependence between ATP production and chromatin motion. Indeed, at the scale of minutes, inhibiting ATP production stalls single loci movement (7). Also, active (ATP-dependent) nuclear processes were shown to drive coherent motion of chromatin at the scale of the entire nucleus (9, 18). By tracking individual loci or by computing the dynamics of chromatin at the scale of the entire nucleus, transcription activation was reported to slow down chromatin motion (8, 15, 19, 20). Thus, it seems that changes in chromatin motion depend on alterations of nuclear processes and activity.

As proliferating cells progress through their cell cycle, they undergo profound changes in nuclear organisation (21, 22) and activities (23). Upon mitotic exit, chromosome decompaction accompanies re-establishment of nuclear architecture at the onset of the G1 growth phase. Duplication of DNA content in S-phase is followed by the G2 growth phase and preparation for cell division.

Using single nucleosome imaging on the time-scale of one second, Iida *et al.,* observed that nucleosome dynamics remain in a steady-state all along the cell cycle (10, 24) and suggested that nucleosome motion is mainly driven by thermal fluctuations in interphase, and is neither affected by the replication nor the doubling of DNA content. Using single particle tracking of telomeres (25), pairs of loci (26) or chromatin domains (27), variations in chromatin dynamics, when monitored on a minute time-scale, were observed during different phases of the cell cycle. These studies clearly suggest that, at a time-scale compatible with common nuclear mechanisms (transcription, repair, rearrangement…), chromatin dynamics vary during the cell cycle, yet the underlying mechanisms remain to be elucidated.

Pursuing the quest of connecting dynamics, structure, and function, we characterised and quantified changes in chromatin dynamics at the scale of the entire nucleus during G1, S and G2 phases. To do so, we adapted the High-resolution Diffusion mapping (Hi-D) technique (15, 28) (see Methods) to precisely analyse recorded motion expressed as mean squared displacement (MSD) of chromatin at nanoscale resolution. To analyse the recorded MSD, we implemented a two-regime model that captures chromatin diffusion at short time-scales and a deterministic drift at longer times, consistent with the action of active forces. Spatially resolved maps of chromatin diffusion regimes within the imaged nucleus were generated.

We show that chromatin motion progressively decreases during S to G2 phases, compared to G1. The measured decrease in dynamics cannot be explained by the activity of DNA polymerases or the progressive entrapment of sister chromatids by cohesive cohesin.

It appears that the doubling of DNA content and the resulting increase in the proportion chromatin occupies within the nuclear volume is responsible for reduced chromatin diffusion observed in G2.

## MATERIALS AND METHODS

### Cell culture

IMR90 human fibroblast cells were grown in complete medium containing DMEM + GlutaMAX (Gibco, 61965-026), supplemented with 15% FBS (Gibco, 10500-064), 1mM sodium pyruvate (Gibco, 11360-039), 1X minimum essential medium non-essential amino acids (Gibco, 11140-035) and 50μg/mL gentamicin solution (Gibco, 15710-049). HeLa sororin-AID, obtained from J.M. Peters (29) and HeLa WT were grown in complete medium containing DMEM + GlutaMAX (Gibco, 61965-026), supplemented with 10% FBS (Gibco, 10500-064), 1mM sodium pyruvate (Gibco, 11360-039), and 50μg/mL gentamicin solution (Gibco, 15710-049). HeLa sororin-AID medium was supplemented with 0.5µg/mL puromycin (Invivogen, ant-pr-1). Cells were cultivated in a 37°C, 5% CO2 incubator and were split before reaching confluency with 0.05% Trypsine-EDTA (Gibco, 25300-054). IMR90 were grown for no more than 14 passages. Cells were tested to be mycoplasma free using MycoAlert mycoplasma detection kit (Lonza, LT07-218).

### DNA plasmids and cell transfection

To assess cell cycle phases in real time the following two constructs were used: for G vs S phases, we used the V40 pNLS-EGFP-L2-PCNA kindly provided by Fabienne Pituello (CBI) and created by Cristina Cardoso’s lab (30). To visualise chromatin, we used a pTRIP-CMV-H2BmCherry created and generously given by Valerie Lobjois and Odile Mondesert (CBI). For G1 vs S/G2 phases, an adaptation of the FUCCI S/G2/M mAG-hGem (MBL Life Science; (31)), created and generously given by Valerie Lobjois and Odile Mondesert, was used. Briefly, the pTRIP-CMV-FUCCI green S-G2-M-P2A-H2BmCherry is composed of the Geminin fused to Azami Green and the H2B fused to mCherry separated by a 2A peptide (P2A). All plasmids were amplified in E.coli DH5α cells and purified using Plasmid Plus Maxi Kit (Qiagen, 12963).

For Hi-D, 4 days before imaging, 150 000 IMR90 cells or 120 000 HeLa (sororin-AID or WT) were plated in glass bottom μ-dish (ibidi, 81156). 3 days before imaging, cells were transfected with pTRIP-CMV Fucci Green S-G2-M-P2A-H2B mCherry (in IMR90 and HeLa, G1 vs S/G2) or doubly transfected with V40 pNLS-EGFP-L2-PCNA and pTRIPCMV-H2BmCherry (in IMR90, G vs S) or solely with pTRIP-CMV-H2BmCherry (in HeLa sororin-AID, G2) with FuGene HD transfection reagent (Promega, E2311). Briefly, 2 μg of plasmids (or 1 μg of each) were diluted in Opti-MEM (Gibco, 51985-026). FuGene reagent was added with a ratio of 4:1 (for IMR90) or 3:1 (for HeLa sororin-AID or WT) for a final volume of 100 μL. The mix was vortexed for 10 seconds and incubated at least 10 min.

Culture medium was changed with fresh complete medium before the addition of the transfection mix and changed again 12h hour after.

### Cell synchronization

To increase the number of cells in S phase before live imaging in the IMR90 population, a thymidine/aphidicolin block was performed as described in (32). 40h before imaging, cells were incubated 14h with 2mM thymidine (Sigma, T1895) then washed three times with PBS at 37°C before addition of fresh complete medium and incubated for 10h. Then 1μg/mL aphidicolin (Sigma, A4487) was added and cells were incubated for 14h. Finally, cells were washed three times with PBS and incubated in medium at least 2 more hours before being processed for live imaging. To synchronise the HeLa sororin-AID in early G2, a similar thymidine/aphidicolin block was performed except that cells were released in the cell cycle 5h before imaging to let them enter into G2, and cells were treated with 500µM of 3-Indoleacetic acid (IAA, Sigma, I5148) to induce sororin degradation 2h before imaging. To synchronise the HeLa sororin-AID in late G2, the thymidine/aphidicolin block is followed, 3h after release, by a 14h incubation with 6µM RO3306 (Invitrogen). Cells were treated with IAA 2h before imaging.

### FACS analysis

To verify the cell cycle distribution in asynchronous IMR90 population, double staining of Hoechst 33324 and Pyronin Y was used (33). One million cells were collected into PBS, fixed for 2h in iced cold 70% ethanol and washed twice with FACS buffer (1× PBS supplemented with 2% FBS and 1 mM EDTA). Cells were incubated in the dark at 37°C, first, for 45min with 2μg/mL Hoechst 33324 (Invitrogen, H3570), second, for 30min with 4μg/mL Pyronin Y (Abcam, ab146350) both diluted in FACS buffer and kept at 4°C in FACS buffer.

CytoFLEX S flow Cytometer was used for the acquisition and CytExpert for the analysis (Beckman Coulter). Hoechst (for DNA) and Pyronin Y (for RNA content) were respectively measured with UV (350nm) and yellow (561nm) lasers. Cells in G0 were identified as the population with 2N DNA content and with a lower RNA content than cells in early S phase.

### Live imaging

On the day of imaging, culture medium is changed to L-15 medium (Gibco, 21083-027) supplemented with 15% FBS for IMR90 or 10% FBS (plus IAA if needed) for HeLa cells. Cells were placed in a 37°C, 5% CO2 humid chamber (OkoLab system). Imaging was performed using a DMi8 inverted automated microscope (Leica Microsystems) featuring a confocal spinning disk unit (CSU-X1-M1N, Yokogawa). A 100x oil immersion objective (Leica HCX-PLAPO) with a 1.4 NA was used for high resolution imaging. Image (16bit) was acquired using Metamorph 7.10.5 software (Molecular Devices) and detected using a CMOS Hamamatsu Flash4 V2+ camera (1003×1433) and 1×1 binning, with sample pixel size of 76 nm. First, sequential acquisition of a multichannel image of the field of interest was done to characterise the cell cycle stage. EGFP (or Azami Green) was imaged using a 488 nm laser with a band pass emission filter of 510-540, mCherry was imaged using a 561 nm laser with a band pass emission filter of 589-625; exposure time was 200ms. Image series of H2B-mCherry, and of PCNA-EGFP, (with corresponding fluorescence filters) of 500 frames, with exposure time of 200ms per frame (5fps) were successively acquired.

### Cell cycle phase determination

For G vs S phases, cells were visually sorted based on the distribution of PCNA staining in the nucleus (diffuse: G, homogeneously punctiform: early S, peripheral punctiform: mid/late S). For G1 vs S-G2, cells were visually sorted based on the presence of Geminin nuclear signal; G1 cells express only H2B-mCherry (Geminin being degraded by the proteasome), while S-G2 cells express both H2B-mCherry and mAG-hGeminin; the specific pattern of H2B in M phase enable elimination of such cells from the analysis.

### Hi-D

#### Pre-processing

Nuclei which undergo deformations, *z*-plan drift or with a low H2B-mCherry intensity (less than twice the intensity of the background) are eliminated as they will bias the detected movements. Bleaching and *XY*-plan drift were corrected using Bleach Correction imageJ plugin (34) and StackReg translation correction imageJ plugin (35) respectively. Finally, to remove high gain noise from timelapse image streams, the Kalman Stack Filter plugin from image J is applied.

#### Optical Flow–Based Reconstruction of Virtual Particle Trajectories

Hi-D analysis is based on the method of (15). Briefly, movements between two consecutive frames were computed by the Farneback Optical Flow based on polynomial expansion (36). This specific method captures the movement at different resolutions using a pyramidal representation, and was the best evaluated among different protocols (19).

Briefly, to track chromatin motion in high-density imaging conditions where single-particle tracking is not feasible, dense Optical Flow was used. Optical Flow estimates frame-to-frame displacement vectors as a velocity field defined at fixed pixel positions (Eulerian description). To obtain trajectory-based information, a set of virtual particles was initialised at pixel centres. Their motion was reconstructed by numerically integrating the velocity field over time (here the field is 11×11 pixels). For each time-step *Δt*, particle positions were updated according to:

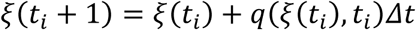

where *q*(*ξ*(*t*_*i*_), *t*_*i*_) denotes the interpolated Optical Flow displacement vector evaluated at the current particle position *ξ*(*t*_*i*_) at time *t*_*i*_. Because the flow field is defined on a regular grid, spatial interpolation was applied at each iteration to evaluate velocities at off-grid particle coordinates. This procedure was repeated sequentially across all acquired image frames to reconstruct virtual particle trajectories at sub-pixel resolution. Importantly, reconstructed trajectories represent local bulk chromatin motion rather than individual molecule trajectories, as each pixel may contain signal from multiple emitters. Quantitative analyses were therefore performed over spatial neighbourhoods rather than interpreting trajectories as single-particle paths.

#### Diffusion characterisation

The MSD is computed for every trajectory as a function of the time interval, *τ*, and the first 20% (100 frames correspond to 20 s) of the experimental MSD was fitted with the following formula that combines anomalous sub-diffusion and drift:

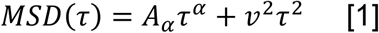

with *α* ≤ 1 so that the anomalous diffusion regime dominates at short times *t* < *t*_*c*_ and that the drift model dominates at longer times *t* > *t*_*c*_, where *t*_*c*_ = (*A*_*α*_⁄*v*^2^)^1⁄(2−*α*)^ is the crossover between the two regimes corresponding to the two terms of the MSD (Equation [1]). Equation [1] combines possibly three dynamical models: free diffusion (*α* = 1), anomalous diffusion (*α* < 1), and directed motion (corresponding to the second term in the rhs. of Equation [1]).

From *A*_*α*_ and *α* obtained from the fitting of the experimental MSD, two independent physical parameters, the diffusion coefficient *D* and the diffusion time *T*_0_ are extracted through the relation 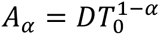. Furthermore, the typical size, *a*, on which the virtual tagged particle freely diffuses at very short times (smaller than *T*_0_) is given by *a*^2^ = *DT*_0_. Each fit is done by minimizing the quadratic error with the experimental MSD (normalised by their maximum values). Only fits with an error inferior to 25% are kept in the following analysis. For a few trajectories, the fitted anomalous exponent *α* systematically reached the upper bound of the allowed interval (0 < *α* < 1.05) and the associated fits were excluded from further analysis. For these few cases, we have preferred to stick with *α* ≤ 1. Indeed, to keep a simple fitting formula, we have assumed that any super-diffusive behaviour is associated with processive motion. A value of *α* larger than 1 would reflect that our simple expression is not perfect and a more elaborate one that smoothly interpolates between the two regimes would be more appropriate. An artificial value of *α* larger than 1 comes from this assumption and is thus assumed to be an unphysical compensation between *α* and *v*.

### Nucleus segmentation

Based on a manually defined intensity threshold for each cell, nucleoli and chromatin ‘holes’ (areas of very low intensity) are segmented and then labelled as nucleoli or holes depending on their features: the nucleolus is usually clearly defined by a high intensity chromatin border in contrast to chromatin holes. These regions with a low signal to noise ratio, as well as, the area outside of the nucleus are excluded from analysis (in white in Figure 2A). Of note, some nuclei may not have a nucleolus in the *z*-plan acquired.

### Single Particle Tracking

PCNA foci are sufficiently spatially separated in space to allow single particle tracking. Detection of foci are performed using the blog_log function from the Skimage library, and trajectories are determined using the library Trackpy on Python (37). Only the detected long trajectories are kept for the MSD analysis (> 300 timepoints). For each nucleus, tracked PCNA foci are classified based on their *MSD*(*τ* = 20*s*) value. Foci in the lowest tercile are classified as foci with slow dynamics, in the medium tercile as foci with medium dynamics and in the highest tercile as foci with high dynamics.

### Simulations

In our Brownian dynamics simulations, the chromatin is modelled as a bead-spring chain composed of *N* = 1000 beads of diameter *a* (with a friction coefficient *ζ* = 3*πηa* where *η* is the solvent viscosity) connected by springs (of equilibrium length *a* and spring constant *k* = 100*k*_*B*_*T*⁄*a*^2^ where *k*_*B*_*T* is the thermal energy). To model the excluded volume interactions between beads, we use the Lennard-Jones potential for the interaction between beads (energy parameter *∊* = 0.0024*k*_*B*_*T* and *σ* = *a*), such that the chain is slightly swollen. Each bead evolves following the overdamped Langevin equation. The bead relaxation diffusion time is defined as *τ*_0_ = *ζa*^2^⁄*k*_*B*_*T* (the bead diffusion coefficient is given by the Einstein relation *D* = *k*_*B*_*T*⁄*ζ*) and a time-step of *δt* = 10^−4^*τ*_0_ is used. Simulation runs were around 10^8^ time-steps (after 10^6^ equilibration time-steps) and averages were done on 25 replicates. The different volume fractions *φ* were obtained by confining the polymer in spheres of radii using 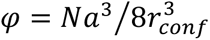. For instance, *φ* = 0.03, 0.13, 0.34 correspond to *r*_*conf*_ = 15.88*a*, *r*_*conf*_ = 9.84*a* and *r*_*conf*_ = 7.14*a*, respectively. 100 beads between the 300 and 400 positions were chosen. The MSDs are computed for each chosen bead, and are then averaged. The simulated MSD were fitted with the following formula:

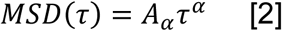

Since, for simplicity, we did not introduce any drift due to the presence of molecular motors in these simulations, we assume *v* = 0 and remove the second term of Equation [1]. The diffusion coefficient *D* was extracted from the simulations similarly to the experimental characterisation of chromatin dynamic parameters. As illustrated in Figure S6C, we clearly observe at very short time-scales, i.e. for lag-times *τ* ≪ *τ*_0_, that the MSD is purely diffusive with *α* = 1, *MSD*(*τ*) = *Dτ*. This is in agreement with polymer theory (38), since at these short lag-times the monomer does not have the time to feel its neighbours and therefore diffuses freely. The relaxation diffusion time can be estimated by the intersection between these two asymptotic regimes leading to 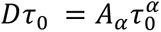 which yields

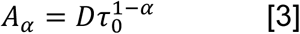

By doing so we confidently find *τ*_0_ which is around 2 or 3 times the value 10^4^*δt* we have chosen in the simulations. However, in experiments, we do not have access to a time-resolution allowing us to observe this diffusive behaviour at short lag-times. So, we propose another way to obtain this value by plotting *Log*(*A*_*α*_) vs. *α* for many numerical replicas and then extracting *D* and *τ*_0_ by assuming Equation [3] (see for instance Figure 1B and 5G). By doing so, we find a value for *τ*_0_, that we noted *T*_0_ as in experiments, which is larger by a factor of approximately 60 (for a fitting window equal to the first 10% of lagtimes) and 120 (for a fitting window of 20%), i.e. *T*_0_ = 6 − 12 × 10^5^ *δt*. This discrepancy comes from the fitting procedure of the MSD for each replica (noise and choice of the fitting window), since with this protocol we infer a characteristic time of diffusion at short times through a fit at long times.

**Figure 1:**
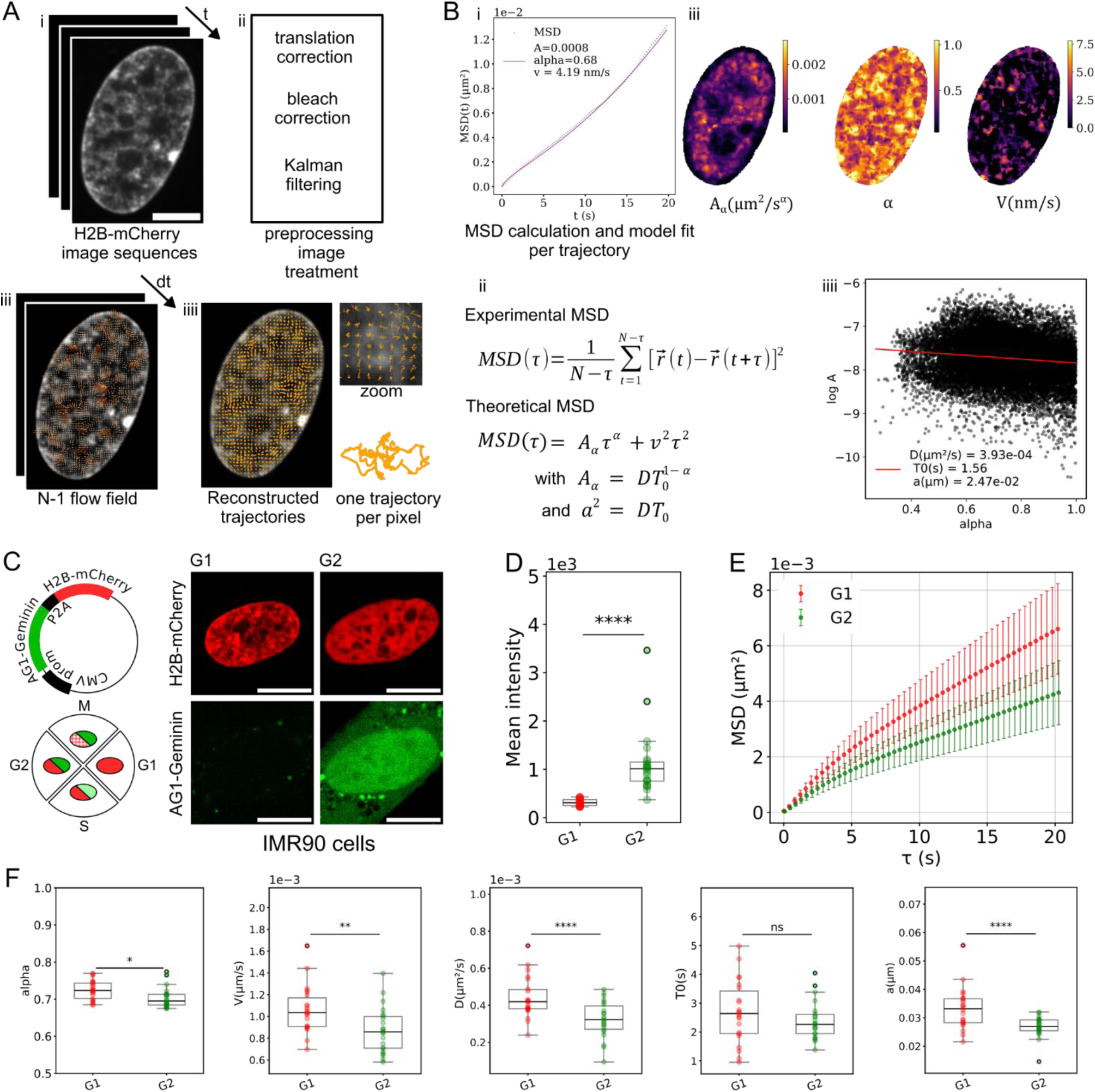
Hi-D–based analysis of chromatin dynamics reveals reduced chromatin mobility during G2. A) Trajectories reconstruction pipeline: i. Image acquisition of a densely fluorescent object (chromatin labelled with H2B-mCherry); ii. Correction and filtering of the image series; iii. Extraction of the flow field between two consecutive images with Farneback dense optical flow; and iv. Trajectory reconstruction, with one trajectory assigned to the initial pixel (for clarity, only a subset of the trajectories is plotted on the image). B) Hi-D analysis and extraction of physical parameters. i and ii. Computation of the experimental MSD for each trajectory fitted with the theoretical anomalous diffusion and drift model. iii. Extraction of the diffusion amplitude (*A*_*α*_), the anomalous exponent *α* and the velocity (*v*) for each trajectory and plotted on the initial pixel. iv. Extraction from *A* and *α* of the diffusion coefficient (*D*), a characteristic time (*T*_0_) and the theoretical size of a particle (*a*), for the analysed region of the nucleus. C) Scheme of the FUCCI plasmid used and the resulting cell cycle distribution of the fluorescent markers. Representative images of G1 and G2 IMR90 nuclei. Scale-bar represents 10 µm. D) Mean intensity of the first frame of the analysed nuclei. Lower and upper hinges correspond to the first and third quartiles, the horizontal line represents the median, whiskers extend to the largest value no further than 1.5 x interquartile range. Dots indicate individual cells. Statistical testing by means of two-tailed Welch’s t test (ns, p<0.05 *, p<0.01 **, p<0.005 ***, p<0.001 ****). E) For cells in G1 (N=22, red) and G2 (N=22, green): mean±SD of the mean MSD of each nucleus. F) Box plots of the physical parameters (*α*, *v*, *D*, *T*_0_ and *a*) extracted from the MSDs shown in Figure 2B for cells in G1 or G2. Box plot description similar to Figure 1D.

## RESULTS

### Chromatin motion decreases between G1 and G2

We have previously shown that global chromatin motion upon exit from quiescence was reduced and correlated with transcription reactivation in G1 (15). We investigated here how cell cycle progression affects chromatin motion.

We characterised chromatin motion at the scale of the entire nucleus using an optimised Hi-D method as described in ((15); and Methods). Hi-D enables analysing the motion of densely distributed molecules such as fluorescently tagged histones incorporated into chromatin (Figure 1A and 1B). Briefly, movements of chromatin labelled with H2B-mCherry are quantitatively retrieved by dense optical flow (36). Images are acquired at 5 fps over 1min 40s (500 frames). By integrating flow fields, a virtual particle trajectory at nanoscale resolution is computed and assigned to the initial pixel of the imaged nucleus, resulting in more than 10,000 trajectories per nucleus (Figure 1A). Of note, some nuclear regions with very weak fluorescence intensity and thus a low signal to noise ratio were eliminated from further analysis. To characterise the motion of the labelled chromatin within flow fields, an experimental MSD is computed from the trajectories. The first 20% (100 frames equal to 20s) of the cumulative MSD data are fitted using an anomalous diffusion and drift model: *MSD*(*τ*) = *A*_*α*_*τ*^*α*^ + *v*^2^*τ*^2^ (Equation [1], Figure 1B). The following physical parameters are extracted: the anomalous exponent *α*, which characterises the diffusion mode of a particle with *α* = 1 corresponding to normal diffusion and *α* < 1 to subdiffusion; the velocity *v*, which characterises the directed motion resulting from active forces, and *A*_*α*_, the diffusion amplitude (µm^2^/s^α^) related to many parameters such as temperature, nucleoplasm viscosity and the size of the tracked particle. From a linear regression of Log(*A*_*α*_) versus the exponent *α* we deduced the effective coefficient of diffusion *D* (in µm^2^/s), the relaxation time *T*_0_ (a characteristic time at which a particle starts to diffuse abnormally) and the effective size of the tracked particle *a* (in µm) (more precisely the variance of the step length distribution), resulting in one unique value of *D*, *T*_0_ and *a* per nucleus (or per segmented area). This model with two regimes allows us to capture the diffusive nature of the chromatin in combination with a deterministic drift motion at long times (generated by active forces).

The fluorescence intensity (correlated with chromatin density), the calculated trajectory and the resulting physical parameters *A*_*α*_, *α*, and *v* are then determined at precise positions within the nucleus. These data enable nanoscale mapping of chromatin dynamics. It is important to underline that using Hi-D, we follow flow fields of fluorescently labelled chromatin fibres as opposed to single nucleosomes. Hence, the diffusion coefficient *D* is based on probing the behaviour of the fibres at a larger length scale than the one of a single monomer as when applying SPT. The deduced size of the fictitious particle *a* corresponds to the correlation length of the concentration field, and is expected to be larger than the nucleosome size.

We labelled IMR90 human fibroblasts with H2B-mCherry (chromatin marker) and mAG-Geminin using an adapted version of the FUCCI system (31) to discriminate cell cycle phases in an asynchronous population. G1 cell nuclei are only stained red and S/G2 cells are stained red and green (Figure 1C). Note that the IMR90 primary human fibroblast cell line used here has a relatively long cell cycle (population doubling in ∼ 3 days) which results in a minority of the cells in S phase at the time of observation (∼ 2%; Figure S1A) ensuring that the vast majority of the green nuclei analysed here are effectively in G2 and not in S phase. Furthermore, we also observed that H2B-mCherry mean intensity of the G2 nuclei is more than two-fold greater than the intensity in G1 nuclei (corresponding to a doubling of the DNA content in our G2 population). The mean area is similar between the two conditions (Figure 1D, Figure S1B). Importantly, there is no significant correlation between the mean MSD and the mean intensity in a given population (in G1 with *R*^2^ = 0.102 or in G2 with *R*^2^ = 0.098), thus ruling out the possibility that the trajectories detected by Hi-D are influenced by H2B-mCherry intensity (Figure S1E).

The mean MSD recorded in G2 is strongly reduced compared to the one measured in G1 (*MSD*(*τ* = 20*s*) = 6.60 ± 1.63 × 10^−3^µm^2^ in G1 and 4.31 ± 1.15 × 10^−3^ µm^2^ in G2 with *p* = 5.84 × 10^−6^ as shown in Figure 1E). Such decrease in diffusion is coherent with reports using SPT methods at a similar temporal scale in different cell types (25, 27). In formaldehyde fixed cells, measured MSD of chromatin is within the technical background, one order of magnitude below the computed MSD in living cells (Figure S1D). We measure a significant decrease in all parameters determined (*v*, *D*, *a*) for chromatin motion of cells in G2 compared to G1 (Figure 1F).

The anomalous exponent *α* of a Gaussian polymer following Rouse dynamics is known to be 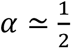 (14). Experimentally, we determined a mean anomalous exponent *α* ≃ 0.71 as shown in Figure 1F. These rather high values of *α* can be explained by the fact that the Hi-D approach consists in tracking a chromatin concentration field, recording the diffusive behaviour of the fibre assimilated to a polymer at a larger length scale than the one of a single nucleosome or small DNA segment. These values are consistent with the sub-diffusive nature of chromatin as previously measured for whole chromatin motion (9, 15). Between G1 and G2, *α* decreases by 0.02 (from 0.725 ± 0.025 in G1 to 0.703 ± 0.027 in G2 with *p* = 1.04 × 10^−2^) (Figure 1F). This small variation from a physical point of view, is not relevant to deduce that the anomalous diffusion regime of chromatin changes, leading us to conclude that the mode of diffusion of the chromatin is not impacted by the cell cycle progression.

We observed that, in live asynchronous populations, chromatin dynamics at the scale of the entire nucleus, is reduced in cells in G2 compared to cells in G1. Major differences in G2 from G1 nuclei are the doubling of the DNA content and the cohesion between the two sister chromatids. We postulate that these events could be responsible for the decrease in chromatin motion.

### Chromatin dynamics are influenced by its position within the nucleus, but the overall changes in motion remain comparable between G1 and G2

If indeed the decrease in chromatin motion in G2 is due to the doubling of the DNA content and the action of cohesive cohesins on sister chromatids, chromatin dynamics are expected to change in all areas of the nucleus.

We, therefore, segmented IMR90 nuclei in distinct regions: the nuclear periphery (NP) corresponding to a rim of 760 nm (10 pixels) at the nuclear border, the nucleolar periphery (NLLP), corresponding to a rim of 760 nm at the nucleolar border, finally the rest which is the nuclear interior (NI) (Figure 2A). Hi-D enables computing chromatin motion originating in each of these regions and mapping the spatial distribution of extracted physical parameters.

**Figure 2:**
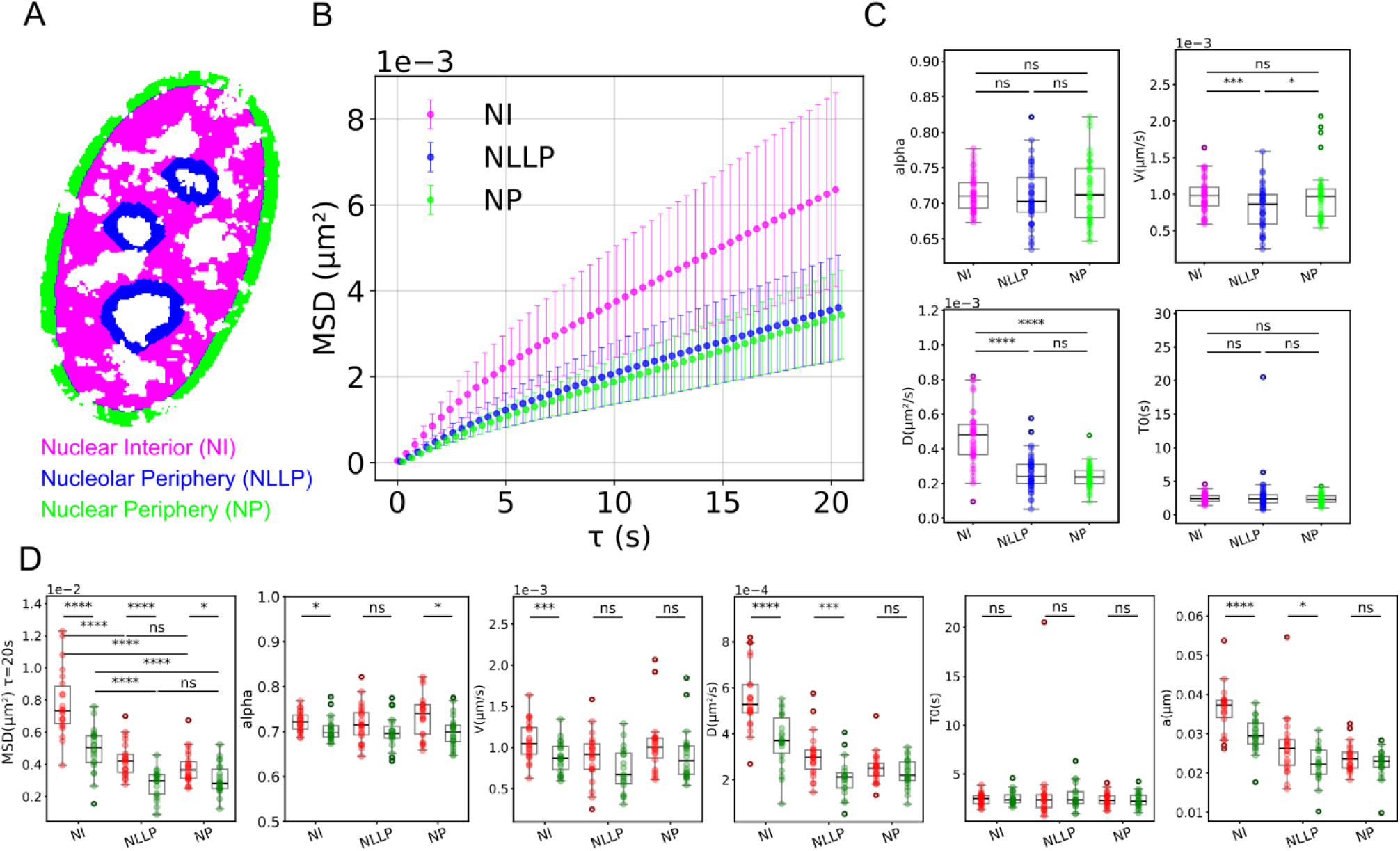
Chromatin dynamics vary according to subnuclear localisation. A) Representative image of a segmented IMR90 nucleus with the nuclear interior (NI), nucleolar periphery (NLLP) and nuclear periphery (NP). White pixels (corresponding to pixels associated with trajectories of displacement > 1 µm, *α* ≥ 1, nucleolar interior and chromatin holes; see Methods section) were eliminated from the analysis. B) For NI, NLLP and NP: mean ± SD of the mean MSD of each segmented nucleus (*N* = 44, same cells as in Figure 1). C) Box plots of the physical parameters extracted from the MSDs shown in panel D (*α*, *v*, *D* and *T*_0_) in the NI, NLLP and NP. Box plot description similar to Figure 1D D) Box plots of the mean *MSD*(*τ* = 20*s*) and the physical parameters (*α*, *v*, *D*, *T*_0_ and *a*) of G1 and G2 nuclei segmented in NI, NLLP and NP. Box plot description similar to Figure 1D. For *α*, *v*, *D*, *T*_0_ and *a*, only the statistical differences between G1 and G2 are shown.

In formaldehyde fixed cells no differences between NP, NLLP and NI were detected (Figure S1D, Figure S2A). We plotted the mean chromatin MSD in each of these regions in living cells. A decrease in motion was measured within both NP and NLLP compared to the nuclear interior: *MSD*(*τ* = 20*s*) = 3.44 ± 1.03 × 10^−3^ µm^2^ for NP, 3.61 ± 1.22 × 10^−3^ µm^2^ for NLLP, and 6.36 ± 2.26 × 10^−3^ µm^2^ for NI (Figure 2B). Chromatin motion is reduced in heterochromatin-rich regions compared to euchromatin-rich ones. *MSD*(*τ* = 20*s*) between G1 and G2 decreased in all of the three identified areas, NI, NP and NLLP (Figure 2D). Thus, the decrease in chromatin motion in G2 quantified by *MSD*(*τ* = 20*s*) seems to be independent of the chromatin localisation (or type, i.e. heterochromatin and euchromatin).

To further investigate this point, we extracted physical parameters from the MSDs (Figure 2B). We found that the diffusion coefficient *D* decreases at the nuclear periphery. This reduction correlates with the physical properties of chromatin in these regions. At the NP, chromatin fibres are known to tether to the nuclear envelope via LADs (39), while in the NLLP, chromatin is compacted by binding proteins, forming heterochromatin regions. Differences in chromatin motion between euchromatin- and heterochromatin-rich regions can also be linked to the parameter *a*, which represents the typical diffusion size of chromatin according to fluctuations in concentration within the flow field. The observed decrease in *a* in heterochromatin-rich areas may be linked to the compaction of the polymer (Figure S2B). The velocity *v*, a parameter related to active processes, appears significantly smaller in the NLLP compared to the NI. The absence of velocity in chromatin fibre motion suggests that there is less, notably transcriptional, activity in the peripheral areas. However, while the NP is also heterochromatin-rich, *v* within the NP and the NI do not vary. This status quo could be due to small and local deformations of the nuclear envelope, sometimes visible in the movies, and that could contribute to greater velocity *v* in this area. Interestingly, when we separate the nuclei in G1 and G2 populations, the decrease observed in (*D*, *a*, *v*) values are stronger in euchromatin-rich regions (NI) compared to heterochromatin-rich regions (NLLP and NP) (Figure 2D). It appears that the phenomenon responsible for the motion decrease in G2 has a greater influence in relatively relaxed regions rather than in already constrained regions of the nucleus. On the other hand, we did not identify any changes in other parameters such as the anomalous exponent *α* and the characteristic diffusion time *T*_0_. This suggests that, at the observed temporal scale, there are no major changes in the nature of the chromatin polymer between eu- and heterochromatin regions.

By leveraging Hi-D dynamics data, we identified differences in chromatin motion as a function of intranuclear spatial localization, thereby confirming that chromatin composition and nuclear architecture influence chromatin motion. We also observed that the reduced chromatin motion in G2 is a global phenomenon and is not restricted to certain areas, even if the dynamical parameters which decrease (*D* or *v*) are not identical in all areas. Our results support that a ubiquitous change in chromatin/ nuclear organisation or conformation between G1 and G2 nuclei is responsible for reduced chromatin dynamics.

### Chromatin motion progressively decreases during S phase

During replication in S phase DNA content is duplicated and cohesive cohesins are loaded on chromatin (40–42). To study S phase nuclei, we transfected IMR90 cells with EGFP-PCNA and H2B-mCherry plasmids (Figure 3A). PCNA, a protein of the replisome recruited at the replication site, forms foci exclusively in S phase (43). The pattern of foci distribution can be used to classify nuclei as they progress through S phase, with foci mainly located in the NI (euchromatin regions) in early S, and in the NP and NLLP (heterochromatin regions) in mid/late S (30, 44) (Figure 3C). To discriminate cells in G1 from cells in G2 (both with a diffuse PCNA staining), we measured the intensity of H2B-mCherry fluorescence (Figure 3B). As shown previously (Figure S1B and S1C), the high mean intensity of nuclei labelled with H2B-mcherry is characteristic of G2 nuclei. Although we measured a small change in surface area of the observed nuclei in S phase compared to G phases, dynamic parameters of the chromatin motion do not correlate with this change (*R*^2^ = 0.004, *p* = 0.87; Pearson, Figure S3A and S3B). Of note, to improve our chances to observe nuclei in S phase in our population (Figure S1A) we performed a mild double block thymidine/aphidicholin and released the cells at least 2h before observation.

**Figure 3:**
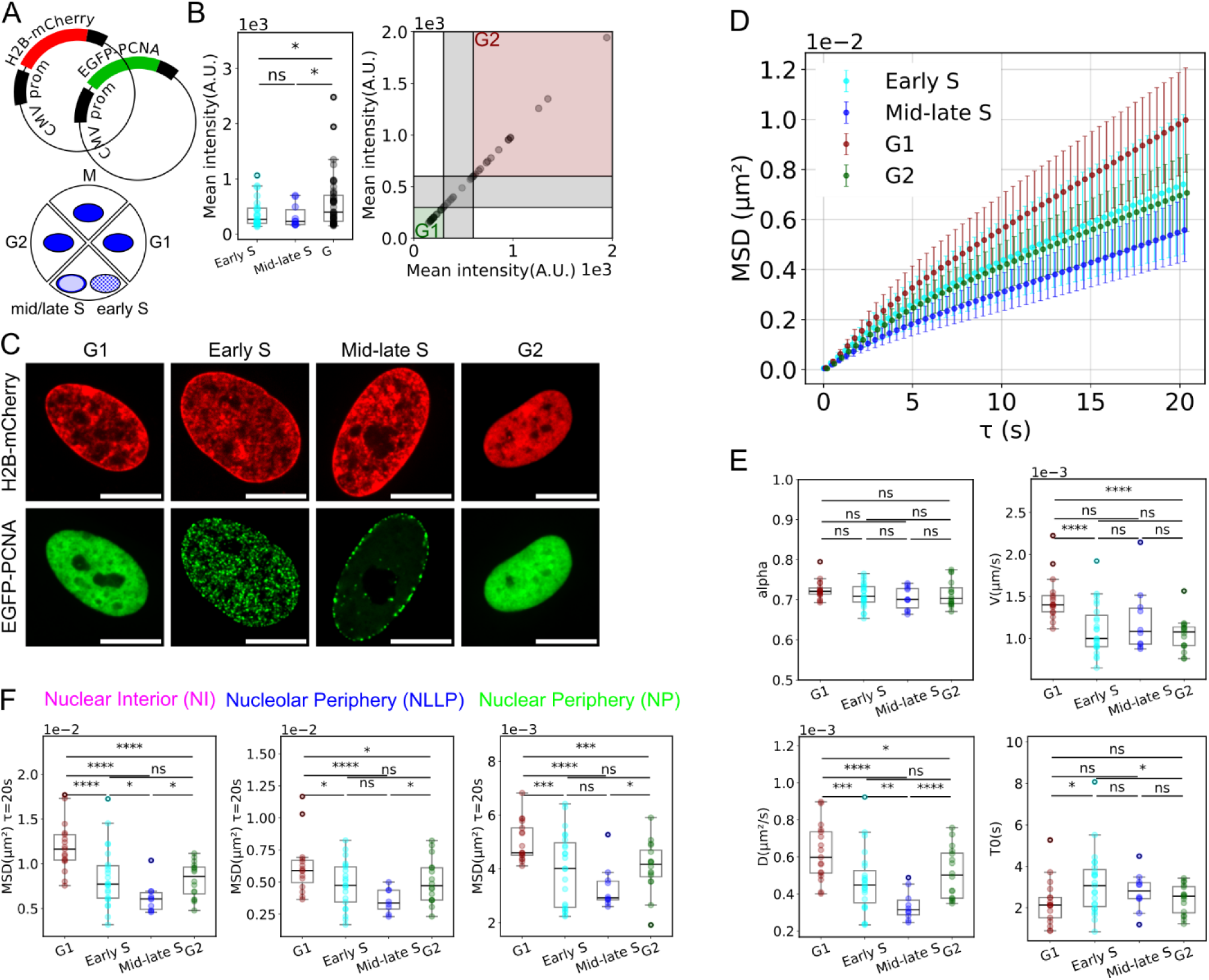
Chromatin dynamics during S phase progression. A) Scheme of the PCNA and H2B plasmids used and the resulting cell cycle distribution of the fluorescent marker (PCNA). B) Box plot depicting the mean intensity of the first frame of the analysed nuclei (only the G phase nuclei, i.e. with a diffuse PCNA staining, are shown). G phase nuclei with an intensity smaller than 300 are associated with G1 (red square) and the ones with an intensity greater than 600 are associated with G2 (green square). C) Representative images of G1, early S, mid/late S and G2 nuclei. Scale-bar represents 10 µm. D) For cells in G1 (N=17, dark red), early S (N=23, cyan), mid/late S (N=10, blue) and G2 (N=16, dark green): mean±SD of the mean MSD of each nucleus. E) Box plots of the physical parameters (*α*, *v*, *D* and *T*_0_) extracted from the MSDs shown in Figure 3D for cells in G1, early S, mid/late S and G2. Box plot description similar to Figure 1D.

The computed *MSD*(*τ* = 20*s*) in G2 nuclei drops significantly by 30% compared to G1 (Figure 3D). Chromatin motion already starts to slow down between G1 and early S (*MSD*(*τ* = 20*s*) = 9.97 ± 2.03 × 10^−3^µm^2^ in G1 and *MSD*(*τ* = 20*s*) = 7.40 ± 2.79 × 10^−3^µm^2^ in early S with *p* = 2.42 × 10^−3^). Motion further decreases when cells are in mid/late S (*MSD*(*τ* = 20*s*) = 5.57 ± 1.25 × 10^−3^µm^2^ in late S with *p* = 6.64 × 10^−7^). Overall, as cells progress through S phase, greater constraints lead to a decrease in diffusive movement. Accordingly, we measured smaller *D* values during S phase (Figure 3E). We also observed a significant decrease in velocity *v* early in S-phase compared to G1. We noticed that chromatin motion slightly increases between mid/late S and G2 (*p* = 1.67 × 10^−2^). We attribute this last difference to the G1 versus G2 discrimination method we employed (based on H2B-mCherry intensity only) leading us to compose with populations that are only enriched in G1 and G2 rather than mutually exclusive when FUCCI is employed.

To go further, *MSD*(*τ* = 20*s*) significantly decreases between G1 and early S in the NI while this decrease is only attenuated in the NLLP and NP. However, when we analysed chromatin motion in mid/late S phase, *MSD*(*τ* = 20*s*) also strongly decreased in the NLLP and NP (Figure 3F, Figure S3D). This phenomenon correlates with the pace of replication: in the NI, constrains arise in early S simultaneously with replication initiation, while in the NP and NLLP, constrains dominate in mid/late S concomitantly to processive replication. These observations strongly support the hypothesis that the decrease in global chromatin motion during the cell cycle arises from an increase in constrains resulting from an increase in chromatin density and/or the entrapment of sister chromatids together by the cohesive cohesins.

To conclude, using spatially resolved dynamic mapping of chromatin motion, we identified a reduction in chromatin motion during S phase progression. Since active processes such as transcription affect chromatin motion (8, 20), one could ask whether DNA replication, through DNA polymerase activities, could alter chromatin motion, at the scale of the entire nucleus.

### Replication / DNA polymerase activity does not modify chromatin motion

To investigate if the replication process *per se* influences chromatin dynamics, we tracked EGFP-PCNA foci in addition to H2B-mcherry (500 frames at 5 frame/s) in S phase cells using SPT (see Methods). The number of PCNA foci in early S phase varies from cell to cell as observed previously (30, 45). We found that the number of replication foci (foci density) and chromatin dynamics within the entire nucleus (characterised by the value of the *MSD*(*τ* = 20*s*)) in our cell population are uncoupled (*R*^2^ = 0.003; *p* = 7.88 × 10^−1^; Pearson, Figure 4A), suggesting that chromatin dynamics are independent of the number of PCNA foci.

**Figure 4:**
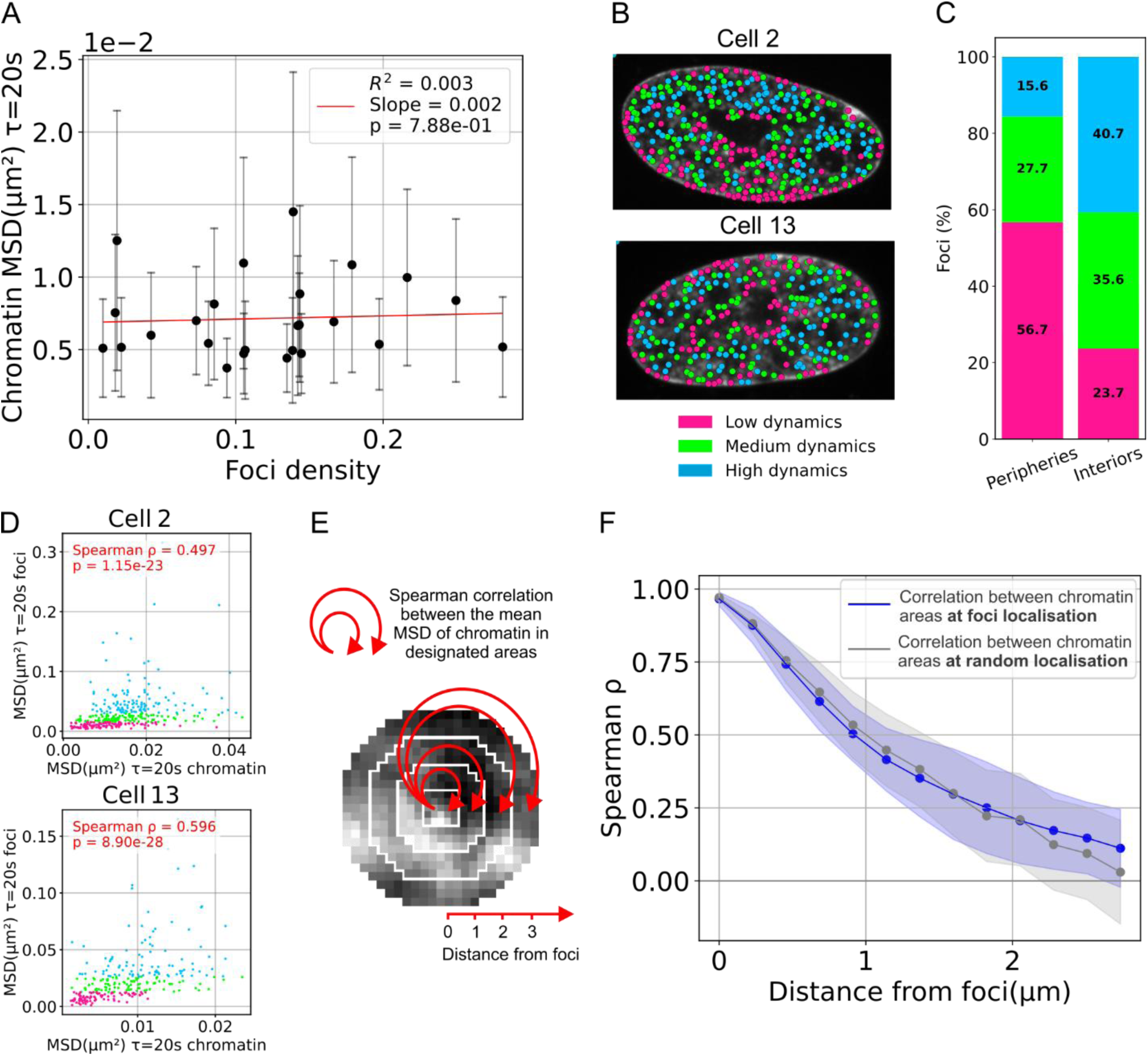
Replication foci dynamics and chromatin motion in S phase. A) Plots for cells in early S and mid/late S phases of the PCNA foci density in nuclei over *MSD*(*τ* = 20*s*) ± *SD* of the chromatin and associated linear regression with p-value from Pearson correlation. B) For 2 nuclei, plot of the tracked PCNA foci color-coded according to their dynamics on the corresponding H2B-mCherry signal in greyscale. Pink, green and blue foci correspond to foci with a *MSD*(*τ* = 20*s*) in the first, second and third tercile respectively of each nucleus (corresponding to a low, medium and high foci dynamics). C) Percentage of foci with a low, medium and high dynamics in the peripheries (NP+NLLP) and in the nuclear interior (NI) with color-codes similar to Figure 4B. D) Scatter plot of the *MSD*(*τ* = 20*s*) of foci versus the *MSD*(*τ* = 20*s*) of the chromatin at the same location, in the two nuclei shown in Figure 1B. Each dot is color-coded according to its dynamic define in Figure 1B. A Spearman correlation is performed and the correlation coefficient *ρ* as well as the p-value is displayed. E) Schematic representation of the method used to plot the Figure 4F. A Spearman correlation is performed between the mean value of the chromatin *MSD*(*τ* = 20*s*) at the center, start of the arrow (a dot of a 3 pixels radius) and the mean value of the chromatin *MSD*(*τ* = 20*s*) of increasing size circle, end of the arrow (with a 3 pixels rim). Distance from foci is the radial distance. F) Plot of the value of the Spearman correlation coefficient *ρ* ± *SD* as a function of the radial distance. In blue, chromatin at a foci localisation in S phase nuclei. In grey, chromatin at random localisation in G phase nuclei.

MSDs of PCNA foci were computed and the value of *MSD*(*τ* = 20*s*) was used as a proxy for their mobility (a high *MSD*(*τ* = 20*s*) corresponds to high foci motion and vice versa). Tracked PCNA foci were classified as low, medium and highly dynamic foci based on their *MSD*(*τ* = 20*s*) value (as shown for two cells, Figure 4B). Only PCNA molecules engaged in DNA replication can be tracked since unbound PCNA molecules diffuse too quickly to be followed by the methodology used here (46). Interestingly, PCNA foci with medium to high dynamics are predominantly present in the NI whereas the slow-moving ones are mostly associated with the NP and the NLLP (Figure 4B and 4C). Hence, PCNA foci dynamics, similar to chromatin dynamics, depend on their nuclear localisation.

At each individual PCNA focus, both motion of the PCNA focus itself as well as motion of the underlying H2B staining can be tracked in the same location successively by SPT and Hi-D, respectively. We observed that slow moving foci are associated with chromatin areas with lower dynamics, and conversely, fast moving foci are associated with chromatin areas with higher dynamics. There is a positive monotonic relationship between the motion of the replication foci and the chromatin at the same location, as shown for two nuclei (Spearman coefficient *ρ* = 0.497; *p* = 1.15 × 10^−23^; *ρ* = 0.596; *p* = 8.90 × 10^−28^; Figure 4D). This raises the question of whether chromatin dynamics drive PCNA foci dynamics, or vice versa.

To tackle this question, we tested if the presence of PCNA foci influences the dynamics of the chromatin underneath. To do so, we measured chromatin dynamics by Hi-D in the area (of 3 pixels radius) in which a PCNA focus is present. We calculated the Spearman correlation coefficient *ρ* between the dynamics of the chromatin located directly at a PCNA focus and the chromatin dynamics at concentric rings of increasing radii around the focus (Figure 4E). We observed that the correlation coefficient decreases with increasing distance from the focus (Figure 4F, blue). When we performed the same analysis in cells in growth phases (G1 or G2) by selecting a random chromatin location we observed the same decrease in the correlation coefficient (Figure 4F grey). This suggests that the presence of a replication focus does not alter the correlation length of the chromatin dynamics.

To sum up, replication foci and chromatin dynamics correlate and, locally, the activities within PCNA loaded replication sites do not seem to perturb chromatin motion in its immediate vicinity. Hence, it appears that, at the spatio-temporal scale observed here, chromatin motion drives motion of the replication foci. Thus, the decrease in chromatin dynamics observed in S phase progression with Hi-D is not due to the loading or activity of the replication machinery but rather to the progression of DNA duplication.

### Doubling of DNA content but not cohesive cohesin is responsible for the decrease in chromatin motion in G2

To verify the role of cohesive cohesin in chromatin motion, we took advantage of an HeLa Kyoto N-terminally-tagged sororin-AID cell line developed and kindly provided by Mitter *et al.*,(29). Sororin is an essential protein of the cohesive cohesin complex preventing its opening by WAPL in S and G2 phases (47, 48). Loss of sororin proteins abolishes chromatid cohesion by the cohesin complex (29, 40).

First, we measured in HeLa WT cells, transfected with our FUCCI system, the influence of cell cycle progression on chromatin dynamics. As seen in fibroblasts, chromatin dynamics decline in G2 cell nuclei compared to G1 cell nuclei. The calculated diffusion coefficients decreased significantly *D* = 5.12 ± 0,511 × 10^−4^ µm^2^/s in G1, and *D* = 4.50 ± 0.809 × 10^−4^ µm^2^/s in G2 with *p* = 4.36 × 10^−2^; as the velocity: *v* = 1.59 ± 0.27 nm/s in G1, and *v* = 1.16 ± 0.152 nm/s in G2, with *p* = 9.03 × 10^−4^ (Figure S4).

Sororin-AID cells were synchronised in S phase using a double block thymidine/ aphidicolin and released into early G2. After at least 2h of auxin treatment GFP-AID-sororin was acutely depleted (Figure 5B). We found that neither chromatin motion between treated and untreated cells (control: *MSD*(*τ* = 20*s*) = 8.14 ± 1.80 × 10^−3^µm^2^ and sororin KD: *MSD*(*τ* = 20*s*) = 7.80 ± 2.11 × 10^−3^ µm^2^ with *p* = 5.14 × 10^−1^) nor the diffusion parameters extracted (*α*, *v*, *D*, *a* and *T*_0_) varied (Figure 5C, 5D and Figure S5C). These results suggest that, in early G2, loss of sister chromatid cohesion does not significantly affect chromatin dynamics. We next synchronised cells in late G2 through addition of RO-3306, an inhibitor of the catalytic activity of human CDK1/cyclin B1 and CDK1/cyclin A complexes (49). At that stage, sororin depletion led to a slight reduction in chromatin motion (control: *MSD*(*τ* = 20*s*) = 8.85 ± 3.24 × 10^−3^µm^2^ and sororin KD: *MSD*(*τ* = 20*s*) = 6.71 ± 2.97 × 10^−3^ µm^2^ with *p* = 1.99 × 10^−3^, Figure S5D, S5F, S5G). Of note, the H2B intensity signal increased in the sororin KD condition versus the control condition in late G2 (but not in early G2) (Figure S5A, S5E). Hence, the reduced motion may likely be due to condensin binding in late G2. Indeed, a recent study reported that depletion of cohesive cohesin in late G2 promotes early recruitment of condensin (50). Condensin binding and subsequent mitotic chromosome condensation is known to hinder chromatin and nucleosome motion (51, 52) and would be coherent with the reduced motion seen upon depletion of cohesive cohesin we measure.

**Figure 5:**
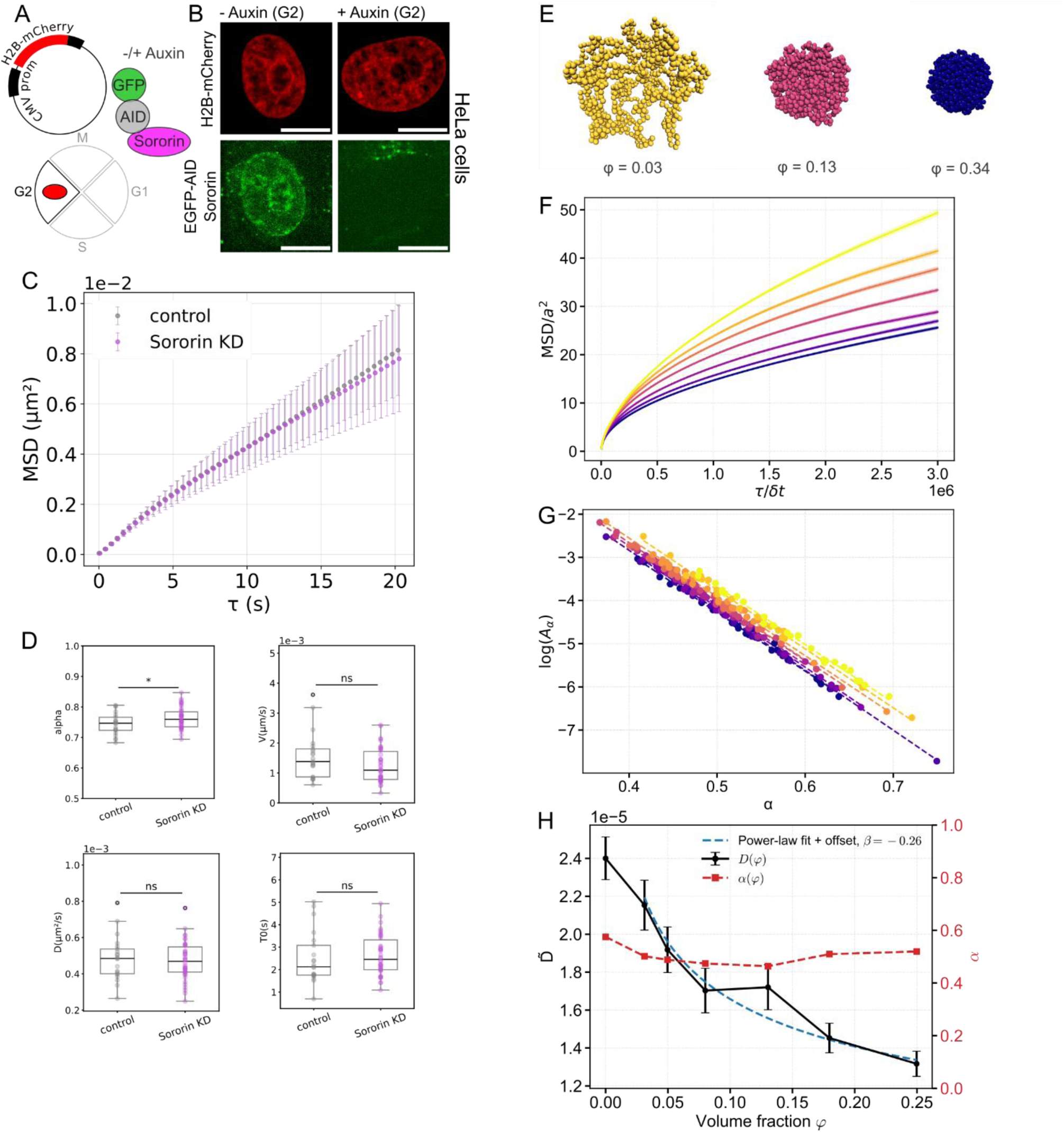
The role of DNA doubling and cohesive cohesins in chromatin dynamics. A) Scheme of the H2B plasmids used, the cell cycle phase studied and the cellular system used (HeLa Kyoto N-terminally-tagged sororin-AID). B) Representative images of HeLa-sororin-AID cells synchronised in early G2 and treated (+ Auxin) or not (- Auxin) to induce the sororin degradation. Scale-bar represents 10 µm. C) For cells, synchronised in early G2, non-treated/control (N=21) and treated with auxin/sororin KD (N=42): mean±SD of the mean MSD of each nucleus. D) Box plots of the physical parameters (*α*, *v*, *D* and *T*_0_) extracted from the MSDs shown in Figure 3D for cells in control and sororin KD condition synchronised in early G2. Box plot description similar to Figure 1D. E) Representative snapshots of chromatin conformation for three confinement radii, *r*_*conf*_ = 15.88, *r*_*conf*_ = 9.84 and *r*_*conf*_ = 7.14, in unit of *a*, corresponding to *φ* = 0.03, *φ* = 0.13 and *φ* = 0.34, using 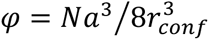 F) Mean squared displacement (MSD) for different volume fractions *φ*, ranging from *φ* = 0 (top, yellow) to *φ* = 0.34 (bottom, purple). Solid curves correspond to ensemble-averaged MSD ± SEM, and dashed lines show power-law fits *MSD* = *A*_*α*_*τ*^*α*^ (Equation [2]). G) Dependence of *Log*(*A*_*α*_) on the exponent *α*. Each point corresponds to an individual simulation seed, with volume fraction increasing from *φ* = 0 (top, yellow) to *φ* = 0.34 (bottom, purple). H) Extracted dynamical parameters from MSD fits: diffusion coefficient 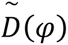 with 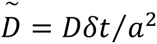 and anomalous exponent *α*(*φ*). The fit of the diffusion coefficient, obtained for intermediate values using a power law, yields an exponent *β* = −0.26 (see Figure S6E).

Experimentally, we observed that sister chromatid cohesion does not impair chromatin motion in early G2, at the observed spatial-temporal scale. To explore in more depth the implication of cohesion on chromatin motion, we turned to polymer simulations.

We performed Brownian dynamics simulations of two polymers, modelled as a flexible bead-spring chain *of N* = 1000 beads with excluded volume interactions (53, 54) (see Methods). These polymers are connected by springs at selected locations along the polymer to simulate attachments due to cohesive cohesins. Four cases with 0,1,2,6 and 10 attachment sites were studied (Figure S6A). For each case, we computed the MSD of 100 beads located between monomer 300 and 400 and averaged them. Sampling was done over 25 replicates. We set the bead size equal to 20 nm (roughly 1 nucleosome plus the linker). In regards to biological data, cohesive cohesins are located at TAD borders so at least every 0.2Mb (29, 41) or every 1000 beads in our model. Therefore, linear density of attachments between our two polymers corresponding to the biological case would be 1 or 2 springs. The recorded MSD changed only slightly between the different cases (Figure S6B), suggesting that cohesion does not have a strong influence on chromatin motion. The strongest effect corresponds obviously to the case of 10 attachments with a decrease of the MSD at 4 × 10^6^ time-steps which is lowered from 60 to 50 (in units of *a*^2^). For only one attachment (close to the biological reality), the MSD of the tracked monomers is weakly impacted only when the attachment is close to them (red case compared to green case, Figure S6A and S6B). These simulations show that sparse attachments between polymers only weakly impair their dynamics, which is in agreement with our experimental observation consecutively to sororin knock-down. We conclude that cohesive cohesins are not responsible for the observed decrease in chromatin motion in G2.

Next, we investigated whether increased confinement restricts chromatin motion by calculating the MSD of confined single polymers. To vary the volume fraction *φ*, we confined the polymer in a sphere of varying radius *r*_*conf*_ by introducing excluded volume interactions with the sphere (see snapshots for various values of *r*_*conf*_ in Figure 5E). Sampling was done over 50 replicates.

We observed that when the volume fraction occupied by the polymer, *φ*, increases, the polymer dynamics, characterised by the computed MSD, decreases (Figure 5F). This decrease is nearly two-fold for small values of *φ* (the MSD at 4 × 10^6^ *δt*). Confinement hence exerts a stronger effect than sister chromatid attachment. We observed no significant change in the value of *α* (*α* = 0.555 ± 0.038, Figure 5H), confirming the experimental results mentioned above. Indeed, for the volume fractions *φ* tested, the Gaussian polymer follows Rouse dynamics with *α* ≃ 1/2. Confinement would have modified the diffusion law to confined diffusion only if the monomers had the time to diffuse over the full confinement sphere radius, i.e. for lag-times larger than *t*_*conf*_ defined by 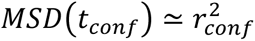. The lag-times studied here (Figure 5F) are much smaller (even in the most restrictive case of *r*_*conf*_ ≃ 8*a*). Hence the diffusion type, and therefore the exponent *α*, does not change.

However, the diffusion coefficient *D* (by using the same protocol as for experimental determination of *D*, see Figure 5G) decreased (Figure 5H). Since the characteristic diffusion time *T*_0_ does not vary with the volume fraction *φ* (Figure S6D), which has also been observed in our experimental data (Figure 1F), it seems that the decrease of *D*(*φ*) is a direct signature of the decrease of the product *DT*_0_. Indeed, 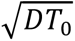 is the characteristic length over which the monomer diffuses freely decreases due to the decrease of the available space for diffusion when *φ* increases. This is also what was observed in experiments (Figure 1F). More quantitatively, we find a scaling law in *DT*_0_ ∝ *φ*^*β*^ with *β* ≈ −0.26 for intermediate values (Figure 5H, Figure S6E). Therefore, doubling the volume fraction leads to a decrease of the MSD at any lag-time by a factor of 2^−*β*^ ≃ 1.2 which is similar to the experimental measurement (Figure 1E).

We conclude that, for a given monomer, the increase of the occupied volume by the polymer directly leads to a larger average number of its neighbouring monomers which directly induces a lower diffusion coefficient *D*(*φ*). The simulated polymer behaviour therefore strongly indicates that the doubling of DNA content in G2, in the nucleus whose volume does not significantly change, is the major contributor to the decrease in chromatin motion observed during the cell cycle.

To conclude, we experimentally tested the implication of cohesive cohesin in chromatin motion and did not observe a significant effect of the removal of cohesion on chromatin dynamics. Polymer modelling revealed that the reduced space available for chromatin motion caused by the doubling of DNA content during S and in G2 phases leads to a marked reduction in chromatin diffusion.

## DISCUSSION

We characterised the evolution of chromatin dynamics over minute-scale intervals throughout interphase progression. We report a progressive decrease in chromatin motion throughout S phase and in G2 that can been observed in all areas of the nucleus, euchromatic and heterochromatic ones. This reduction in chromatin motion, thanks to polymer modelling, reflects confinement due to doubling of DNA content which induces a decrease of the diffusion coefficient, rather than sister chromatid cohesion.

We developed an optimised Hi-D pipeline to track motion of bulk chromatin in specific subzones of a given nucleus at nanoscale resolution. Notably, motion in peripheral regions (of the nucleus and the nucleolus) is more constrained than in the nuclear interior. Since chromatin located at the nuclear rim or the nucleolar periphery are known to be heterochromatin-rich, we conclude that the degree of chromatin compaction influences chromatin motion in agreement with recent work studying chromatin motion at the single nucleosome resolution (10, 11) and original studies using SPT in particular of telomeres (55, 56).

Our results on the motion of chromatin within the entire nucleus provide detailed and quantitative analysis of the dynamic constraints associated with cell cycle progression. This work extends and generalises the observations of earlier work studying specific single chromatin loci, on the increase of constraint in G2. Using single-particle tracking of paired loci, Ma *et al.* (26) showed that following a relaxation phase after mitotic exit in early G1, chromatin progressively condenses from early to late S phase, concomitant with a reduction in mobility. Similarly, Naor *et al.*, (25) using 3D tracking of telomeres, reported that telomere diffusion is cell-cycle dependent, with increased constraints during S and G2 phases relative to G1. By tracking chromatin motion throughout the entire nucleus, we generalise these observations to the entire nuclear space and propose that a ubiquitous process initiated in S phase progressively constrains chromatin dynamics.

Cohesive cohesins, which entrap sister chromatids to ensure proper segregation in mitosis, could be responsible for this decrease in motion. Our experiments and simulations show that cohesive cohesins in early G2 do not impair chromatin dynamics. This could be explained by the stochastic association of these anchors along the genome and their sparse distribution. Cohesive cohesins are not evenly distributed along the chromatin but rather accumulate at TAD boundaries (every 850kb on average) (29). Also, cohesive cohesin loading is not persistent but variable (57). The sheer number of entrapment sites and their variability is insufficient to alter diffusion of chromatin at the minute time-scale.

Our simulations show that the decrease of the MSD due to the presence of attachments is so light that it might not be detectable experimentally.

Alternatively, the duplication of the genetic content in G2 phase in a relatively constant nuclear volume, reducing the available space for diffusion, causes an increase in constraints on chromatin dynamics. Using simulations, the relative decrease of polymer diffusion is around 16% for a doubling of the chromatin volume fraction. This corresponds, albeit slightly smaller, to experimental measurements with a decrease of *D* of 26% between G1 (*D* = 4.39 ± 1.08 × 10^−4^) and G2 (*D* = 3.23 ± 0,97 × 10^−4^) in IMR90 cells (Figure 1F). In fibroblasts, if we assume that a nucleosome is a cylinder (of 10 nm in diameter and 6 nm in thickness), in G1, diploid fibroblasts have 3 × 10^7^ nucleosomes (6 × 10^9^ bp / 200 bp of nucleosome spacing), thus the chromatin of the entire genome occupies ≃ 14 µm^3^. If we consider that a fibroblast has a nuclear volume between 200 µm^3^ and 500 µm^3^, we note a volume fraction occupancy *φ* between ∼3% and ∼7% in G1.

According to our simulations, this small occupancy already imposes constraint on polymer diffusion (Figure 5H). In G2, even though we did not observe significant change by looking solely at the area, it has been reported in the literature that the nuclear volume increases (58, 59). Would it be sufficient to compensate the doubling of DNA, resulting in a constant volume fraction of chromatin along the cell cycle? This has not been formally observed but a body of evidence suggests that the volume fraction occupied by chromatin increases in G2 nuclei. First, by fluorescent microscopy, chromatin in G2 (Figure 1C, Figure S4C) appears denser, with less darker spaces in our cellular models (IMR90 and HeLa cells). In accordance, analysis of the genome fractality by microscopy reveals an increase in chromatin condensation starting from S phase (22). Second, HiC analysis reported an increase in AA and BB compartment frequency of interactions in G2 suggesting that they may be in closer proximity (21). Therefore, it appears that chromatin occupancy increases in G2 compared to G1, and even if it is not a two-fold increase, it would lead to a decrease in diffusion coefficient and a decrease in chromatin dynamics in G2. A similar relationship between nuclear size and chromatin motion was reported in nuclei of *C.elegans* cells during early embryonic development (60, 61).

Regarding the replication phase of the cell cycle, PCNA foci and chromatin dynamics are coupled. Replication foci are more mobile in euchromatic than heterochromatic regions. At the spatio-temporal resolution studied here, the presence of replication foci does not seem to influence the motion of its surrounding chromatin. Importantly, because tracking chromatin motion by HiD relies on the tracking of an intensity field rather than points, we cannot exclude that DNA polymerase activity affects chromatin motion at other scales.

At the scale of the entire nucleus, by computing differentially the sub-S phases (early and mid/late S), we measure a progressive decrease of chromatin motion that coincides with the replication pace. This phenomenon cannot be attributed to the occupancy of the replication machinery on the fibre *per se* but it could be attributed to the progressive increase in chromatin occupancy in the nuclear space. Contrary to human cells, in budding yeast, the nucleus doubles in volume during replication (62). By modelling chromatin doubling during S phase progression in a yeast nucleus, D’Asaro *et al.*, (63, 64) also identified that the presence of catenated chains produced by the replication process lowers the mobility of individual loci. Hence, it is possible that the intertwining of newly replicated chromatin fragments impacts chromatin motion in human cells. Overall, during cell cycle progression, progressive synthesis of chromatin modifies chromatin motion.

Nuclear lamins provide mechanical stability, can control nuclear size (65) and may restrict expansion of the nucleus despite doubling of DNA content. In lamin A mutants, chromatin dynamics increase (66), which is thought to be due to loss of NP chromatin contacts with the envelope. Interestingly, depletion of lamin A alters telomere dynamics along the cell cycle (25). The change in nuclear volume and resulting chromatin occupancy was not envisioned in this study. In mammalian cells, chromatin must remain cohesive to enable condensation at the onset of mitosis when lamins disassemble. This reduced amplitude of motion in G2 may help preserve genome integrity.

In G2, the diffusion coefficient *D* decreases due to the doubling of DNA, but the velocity *v* also decreased. A velocity may originate through a force driving the chromatin polymer in a given direction. We hypothesise that, in chromatin, such movement may be due to the action of molecular motors such as polymerases or loop extruding factors. Note that we measure values on the order of 1 nm/s which is the order of magnitude of molecular motor velocities (around 5 kbp/min for polymerase progression along DNA). In interphase, loop extruding cohesins were shown to provide fast motor activity in vitro (67) and in vivo (68). Interestingly, in G2, the activity of loop extruding cohesins seems to be reduced (observed notably by the stability of the insulation score between TAD in G2 compare to G1 (21)).

This effect could be a rationale for the decrease of *v* observed in G2. Regarding transcription, the global production of mRNA is constant between G1 and G2 despite the doubling of DNA content. To compensate the change in DNA copy numbers, cells in G2 exhibit a decrease in transcriptional burst frequency (69). There is “less” transcriptional activity for a given time period in G2. This phenomenon may further dampen velocity in G2.

Together, variations in chromatin motion occur during different cell cycle phases, as shown in this study during interphase, but culminate in the condensed mitotic chromosome (52). It raises the question on how chromatin motion behaves prior to and following dramatic mitotic chromosome condensation and de-condensation and the associated disorganisation and reorganisation of the chromatin architecture (70) to ensure truthful transmission of the genetic information.

## DATA AVAILABILITY

All datasets obtained during the course of this study are available upon request. We utilised established algorithms for image and statistical analysis as described in the Methods section, coded in Python (version 3.12). The optimised Hi-D pipeline and the homemade Brownian Dynamics codes can be shared upon request.

## SUPPLEMENTARY DATA

Supplementary Data are available online.

## AUTHOR CONTRIBUTIONS

M.RM: conceptualisation, methodology, investigation, formal analysis, supervision, validation, visualisation, funding acquisition, writing-original draft. L.C: conceptualisation, methodology, formal analysis, data curation, investigation, software, validation, visualisation, writing-review. E.LF: investigation, formal analysis, coding H.A: investigation. Q.S: software. M.M: conceptualisation, resources, project administration, supervision, funding acquisition, writing-review. K.B: conceptualisation, resources, project administration, supervision, funding acquisition, writing-review.

## ACKNOWLEDGEMENTS

We thank Odile Mondesert and Valerie Lobjois for sharing FUCCI constructs and Claudia Blaukopf and Jan-Michael Peters for sending us their HeLa Kyoto N-terminally-tagged sororin-AID cell line. We thank the Light Imaging Toulouse CBI (LITC; IBISA and FBI) facility and their expert staff, in particular Vanessa Dougados. We thank all the members of the “Chromatin and Gene Expression” team for insightful discussions as well as Frédéric Beckouët and Wylie Ahmed for critical reading of the manuscript.

## REFERENCES

1. Bonev, B. and Cavalli, G. (2016) Organization and function of the 3D genome. Nat Rev Genet, 17, 661–678.

2. Misteli, T. (2020) The Self-Organizing Genome: Principles of Genome Architecture and Function. Cell, 183, 28–45.

3. Sexton, T. and Yaffe, E. (2015) Chromosome Folding: Driver or Passenger of Epigenetic State? Cold Spring Harb Perspect Biol, 7, a018721.

4. Lieberman-Aiden, E., Berkum, N.L. van, Williams, L., Imakaev, M., Ragoczy, T., Telling, A., Amit, I., Lajoie, B.R., Sabo, P.J., Dorschner, M.O., et al. (2009) Comprehensive Mapping of Long-Range Interactions Reveals Folding Principles of the Human Genome. Science, 326, 289–293.

5. Rao, S.S.P., Huntley, M.H., Durand, N.C., Stamenova, E.K., Bochkov, I.D., Robinson, J.T., Sanborn, A.L., Machol, I., Omer, A.D., Lander, E.S., et al. (2014) A 3D Map of the Human Genome at Kilobase Resolution Reveals Principles of Chromatin Looping. Cell, 159, 1665– 1680.

6. Marshall, W.F., Straight, A., Marko, .F., Swedlow, J., Dernburg, A., Belmont, A., Murray, A.W., Agard, D.A. and Sedat, J.W. (1997) Interphase chromosomes undergo constrained diffusional motion in living cells. Current Biology, 7, 930–939.

7. Heun, P., Laroche, T., Shimada, K., Furrer, P. and Gasser, S.M. (2001) Chromosome dynamics in the yeast interphase nucleus. Science, 294, 2181–2186.

8. Germier, T., Kocanova, S., Walther, N., Bancaud, A., Shaban, H.A., Sellou, H., Politi, A.Z., Ellenberg, J., Gallardo, F. and Bystricky, K. (2017) Real-Time Imaging of a Single Gene Reveals Transcription-Initiated Local Confinement. Biophysical Journal, 113, 1383–1394.

9. Zidovska, A., Weitz, D.A. and Mitchison, T.J. (2013) Micron-scale coherence in interphase chromatin dynamics. Proceedings of the National Academy of Sciences, 110, 15555–15560.

10. Nozaki, T., Imai, R., Tanbo, M., Nagashima, R., Tamura, S., Tani, T., Joti, Y., Tomita, M., Hibino, K., Kanemaki, M.T., et al. (2017) Dynamic Organization of Chromatin Domains Revealed by Super-Resolution Live-Cell Imaging. Molecular Cell, 67, 282–293.e7.

11. Daugird, T.A., Shi, Y., Holland, K.L., Rostamian, H., Liu, Z., Lavis, L.D., Rodriguez, J., Strahl, B.D. and Legant, W.R. (2024) Correlative single molecule lattice light sheet imaging reveals the dynamic relationship between nucleosomes and the local chromatin environment. Nat Commun, 15, 4178.

12. Dutta, S., Ghosh, A., Boettiger, A.N. and Spakowitz, A.J. (2023) Leveraging polymer modeling to reconstruct chromatin connectivity from live images. Biophys J, 122, 3532–3540.

13. Hajjoul, H., Mathon, J., Ranchon, H., Goiffon, I., Mozziconacci, J., Albert, B., Carrivain, P., Victor, J.-M., Gadal, O., Bystricky, K., et al. (2013) High-throughput chromatin motion tracking in living yeast reveals the flexibility of the fiber throughout the genome. Genome Res., 23, 1829–1838.

14. Doi, M. and Edwards, S.F. (1988) The Theory of Polymer Dynamics Oxford University Press, Oxford, New York.

15. Shaban, H.A., Barth, R., Recoules, L. and Bystricky, K. (2020) Hi-D: nanoscale mapping of nuclear dynamics in single living cells. Genome Biology, 21, 95.

16. Tan, T., Wu, J., Si, C., Dai, S., Zhang, Y., Sun, N., Zhang, E., Shao, H., Si, W., Yang, P., et al. (2021) Chimeric contribution of human extended pluripotent stem cells to monkey embryos ex vivo. Cell, 184, 2020–2032.e14.

17. Kocanova, S., Raynal, F., Goiffon, I., Oksuz, B.A., Baú, D., Kamgoué, A., Cantaloube, S., Zhan, Y., Lajoie, B., Marti-Renom, M.A., et al. (2024) Enhancer-driven 3D chromatin domain folding modulates transcription in human mammary tumor cells. Life Science Alliance, 7.

18. Saintillan, D., Shelley, M.J. and Zidovska, A. (2018) Extensile motor activity drives coherent motions in a model of interphase chromatin. Proceedings of the National Academy of Sciences, 115, 11442–11447.

19. Shaban, H.A., Barth, R. and Bystricky, K. (2018) Formation of correlated chromatin domains at nanoscale dynamic resolution during transcription. Nucleic Acids Research, 46, e77.

20. Chu, F.-Y., Clavijo, A.S., Lee, S. and Zidovska, A. (2024) Transcription-dependent mobility of single genes and genome-wide motions in live human cells. Nat Commun, 15, 8879.

21. Nagano, T., Lubling, Y., Várnai, C., Dudley, C., Leung, W., Baran, Y., Mendelson Cohen, N., Wingett, S., Fraser, P. and Tanay, A. (2017) Cell-cycle dynamics of chromosomal organization at single-cell resolution. Nature, 547, 61–67.

22. Lee, S., Liu, X., Ziabkin, I. and Zidovska, A. (2025) Image-based analysis of the genome’s fractality during the cell cycle. Biophysical Journal, 0.

23. Delgado-Román, I. and Muñoz-Centeno, M.C. (2021) Coupling Between Cell Cycle Progression and the Nuclear RNA Polymerases System. Front. Mol. Biosci., 8.

24. Iida, S., Shinkai, S., Itoh, Y., Tamura, S., Kanemaki, M.T., Onami, S. and Maeshima, K. (2022) Single-nucleosome imaging reveals steady-state motion of interphase chromatin in living human cells. Science Advances, 8, eabn5626.

25. Naor, T., Nogin, Y., Nehme, E., Ferdman, B., Weiss, L.E., Alalouf, O. and Shechtman, Y. (2022) Quantifying cell-cycle-dependent chromatin dynamics during interphase by live 3D tracking. iScience, 25, 104197.

26. Ma, H., Tu, L.-C., Chung, Y.-C., Naseri, A., Grunwald, D., Zhang, S. and Pederson, T. (2019) Cell cycle– and genomic distance–dependent dynamics of a discrete chromosomal region. Journal of Cell Biology, 218, 1467–1477.

27. Pabba, M.K., Ritter, C., Chagin, V.O., Meyer, J., Celikay, K., Stear, J.H., Loerke, D., Kolobynina, K., Prorok, P., Schmid, A.K., et al. (2023) Replisome loading reduces chromatin motion independent of DNA synthesis. eLife, 12.

28. Valades-Cruz, C.A., Barth, R., Abdellah, M. and Shaban, H.A. (2024) Genome-wide analysis of the biophysical properties of chromatin and nuclear proteins in living cells with Hi-D. Nat Protoc, 10.1038/s41596-024-01038-3.

29. Mitter, M., Gasser, C., Takacs, Z., Langer, C.C.H., Tang, W., Jessberger, G., Beales, C.T., Neuner, E., Ameres, S.L., Peters, J.-M., et al. (2020) Conformation of sister chromatids in the replicated human genome. Nature, 586, 139–144.

30. Leonhardt, H., Rahn, H.-P., Weinzierl, P., Sporbert, A., Cremer, T., Zink, D. and Cardoso, M.C. (2000) Dynamics of DNA Replication Factories in Living Cells. Journal of Cell Biology, 149, 271–280.

31. Sakaue-Sawano, A., Kurokawa, H., Morimura, T., Hanyu, A., Hama, H., Osawa, H., Kashiwagi, S., Fukami, K., Miyata, T., Miyoshi, H., et al. (2008) Visualizing spatiotemporal dynamics of multicellular cell-cycle progression. Cell, 132, 487–498.

32. Crabbe, L., Verdun, R.E., Haggblom, C.I. and Karlseder, J. (2004) Defective Telomere Lagging Strand Synthesis in Cells Lacking WRN Helicase Activity. Science, 306, 1951–1953.

33. Eddaoudi, A., Canning, S.L. and Kato, I. (2018) Flow Cytometric Detection of G0 in Live Cells by Hoechst 33342 and Pyronin Y Staining. Methods Mol Biol, 1686, 49–57.

34. Miura, K. (2020) Bleach correction ImageJ plugin for compensating the photobleaching of time-lapse sequences. 10.12688/f1000research.27171.1.

35. Thevenaz, P., Ruttimann, U.E. and Unser, M. (1998) A pyramid approach to subpixel registration based on intensity. IEEE Transactions on Image Processing, 7, 27–41.

36. Farnebäck, G. (2003) Two-Frame Motion Estimation Based on Polynomial Expansion. In Bigun, J., Gustavsson, T. (eds), Image Analysis. Springer, Berlin, Heidelberg, pp. 363–370.

37. Crocker, J.C. and Grier, D.G. (1996) Methods of Digital Video Microscopy for Colloidal Studies. Journal of Colloid and Interface Science, 179, 298–310.

38. Rubinstein, M., Colby, R.H., Rubinstein, M. and Colby, R.H. (2003) Polymer Physics Oxford University Press, Oxford, New York.

39. Briand, N. and Collas, P. (2020) Lamina-associated domains: peripheral matters and internal affairs. Genome Biology, 21, 85.

40. Rankin, S., Ayad, N.G. and Kirschner, M.W. (2005) Sororin, a Substrate of the Anaphase-Promoting Complex, Is Required for Sister Chromatid Cohesion in Vertebrates. Molecular Cell, 18, 185–200.

41. Ochs, F., Green, C., Szczurek, A.T., Pytowski, L., Kolesnikova, S., Brown, J., Gerlich, D.W., Buckle, V., Schermelleh, L. and Nasmyth, K.A. (2024) Sister chromatid cohesion is mediated by individual cohesin complexes. Science, 383, 1122–1130.

42. Murayama, Y., Samora, C.P., Kurokawa, Y., Iwasaki, H. and Uhlmann, F. (2018) Establishment of DNA-DNA Interactions by the Cohesin Ring. Cell, 172, 465–477.e15.

43. Prelich, G., Kostura, M., Marshak, D.R., Mathews, M.B. and Stillman, B. (1987) The cell-cycle regulated proliferating cell nuclear antigen is required for SV40 DNA replication in vitro. Nature, 326, 471–475.

44. Celis, J.E. and Celis, A. (1985) Cell cycle-dependent variations in the distribution of the nuclear protein cyclin proliferating cell nuclear antigen in cultured cells: subdivision of S phase. Proc Natl Acad Sci U S A, 82, 3262–3266.

45. Chagin, V.O., Casas-Delucchi, C.S., Reinhart, M., Schermelleh, L., Markaki, Y., Maiser, A., Bolius, J.J., Bensimon, A., Fillies, M., Domaing, P., et al. (2016) 4D Visualization of replication foci in mammalian cells corresponding to individual replicons. Nat Commun, 7, 11231.

46. Zessin, P.J.M., Sporbert, A. and Heilemann, M. (2016) PCNA appears in two populations of slow and fast diffusion with a constant ratio throughout S-phase in replicating mammalian cells. Sci Rep, 6, 18779.

47. Ladurner, R., Kreidl, E., Ivanov, M.P., Ekker, H., Idarraga-Amado, M.H., Busslinger, G.A., Wutz, G., Cisneros, D.A. and Peters, J. (2016) Sororin actively maintains sister chromatid cohesion. The EMBO Journal, 35, 635–653.

48. Prusén Mota, I., Galova, M., Schleiffer, A., Nguyen, T.-T., Kovacikova, I., Farias Saad, C., Litos, G., Nishiyama, T., Gregan, J., Peters, J.-M., et al. (2024) Sororin is an evolutionary conserved antagonist of WAPL. Nat Commun, 15, 4729.

49. Vassilev, L.T., Tovar, C., Chen, S., Knezevic, D., Zhao, X., Sun, H., Heimbrook, D.C. and Chen, L. (2006) Selective small-molecule inhibitor reveals critical mitotic functions of human CDK1. Proceedings of the National Academy of Sciences, 103, 10660–10665.

50. Choi, E.-H. and Kim, K.P. (2025) Cohesin and condensin regulate chromosome topology and play an essential role in maintaining pluripotency in embryonic stem cells. Sci Rep, 15, 9918.

51. Bailey, M.L.P., Surovtsev, I., Williams, J.F., Yan, H., Yuan, T., Li, K., Duseau, K., Mochrie, S.G.J. and King, M.C. (2023) Loops and the activity of loop extrusion factors constrain chromatin dynamics. MBoC, 34, ar78.

52. Hibino, K., Sakai, Y., Tamura, S., Takagi, M., Minami, K., Natsume, T., Shimazoe, M.A., Kanemaki, M.T., Imamoto, N. and Maeshima, K. (2024) Single-nucleosome imaging unveils that condensins and nucleosome–nucleosome interactions differentially constrain chromatin to organize mitotic chromosomes. Nat Commun, 15, 7152.

53. Manghi, M., Tardin, C., Baglio, J., Rousseau, P., Salomé, L. and Destainville, N. (2010) Probing DNA conformational changes with high temporal resolution by tethered particle motion. Phys Biol, 7, 046003.

54. Dasanna, A.K., Destainville, N., Palmeri, J. and Manghi, M. (2013) Slow closure of denaturation bubbles in DNA: twist matters. Phys Rev E Stat Nonlin Soft Matter Phys, 87, 052703.

55. Bystricky, K., Laroche, T., van Houwe, G., Blaszczyk, M. and Gasser, S.M. (2005) Chromosome looping in yeast: telomere pairing and coordinated movement reflect anchoring efficiency and territorial organization. J Cell Biol, 168, 375–387.

56. Bronstein, I., Israel, Y., Kepten, E., Mai, S., Shav-Tal, Y., Barkai, E. and Garini, Y. (2009) Transient anomalous diffusion of telomeres in the nucleus of mammalian cells. Phys Rev Lett, 103, 018102.

57. Stanyte, R., Nuebler, J., Blaukopf, C., Hoefler, R., Stocsits, R., Peters, J.-M. and Gerlich, D.W. (2018) Dynamics of sister chromatid resolution during cell cycle progression. J Cell Biol, 217, 1985–2004.

58. Pennacchio, F.A., Poli, A., Pramotton, F.M., Lavore, S., Rancati, I., Cinquanta, M., Vorselen, D., Prina, E., Romano, O.M., Ferrari, A., et al. (2024) N2FXm, a method for joint nuclear and cytoplasmic volume measurements, unravels the osmo-mechanical regulation of nuclear volume in mammalian cells. Nat Commun, 15, 1070.

59. Fidorra, J., Mielke, T., Booz, J. and Feinendegen, L.E. (1981) Cellular and nuclear volume of human cells during the cell cycle. Radiat Environ Biophys, 19, 205–214.

60. Arai, R., Sugawara, T., Sato, Y., Minakuchi, Y., Toyoda, A., Nabeshima, K., Kimura, H. and Kimura, A. (2017) Reduction in chromosome mobility accompanies nuclear organization during early embryogenesis in Caenorhabditis elegans. Sci Rep, 7, 3631.

61. Yesbolatova, A.K., Arai, R., Sakaue, T. and Kimura, A. (2022) Formulation of Chromatin Mobility as a Function of Nuclear Size during C. elegans Embryogenesis Using Polymer Physics Theories. Phys Rev Lett, 128, 178101.

62. Neumann, F.R. and Nurse, P. (2007) Nuclear size control in fission yeast. J Cell Biol, 179, 593– 600.

63. D’Asaro, D., Arbona, J.-M., Piveteau, V., Piazza, A., Vaillant, C. and Jost, D. (2025) Genome-wide modeling of DNA replication in space and time confirms the emergence of replication specific patterns in vivo in eukaryotes. Genome Biol, 26, 431.

64. D’Asaro, D., Tortora, M.M.C., Vaillant, C., Arbona, J.-M. and Jost, D. (2024) DNA Replication and Polymer Chain Duplication Reshape the Genome in Space and Time. Phys. Rev. X, 14, 041020.

65. Jevtić, P., Edens, L.J., Li, X., Nguyen, T., Chen, P. and Levy, D.L. (2015) Concentration-dependent Effects of Nuclear Lamins on Nuclear Size in *Xenopus* and Mammalian Cells*. Journal of Biological Chemistry, 290, 27557–27571.

66. Bronshtein, I., Kepten, E., Kanter, I., Berezin, S., Lindner, M., Redwood, A.B., Mai, S., Gonzalo, S., Foisner, R., Shav-Tal, Y., et al. (2015) Loss of lamin A function increases chromatin dynamics in the nuclear interior. Nat Commun, 6, 8044.

67. Davidson, I.F., Bauer, B., Goetz, D., Tang, W., Wutz, G. and Peters, J.-M. (2019) DNA loop extrusion by human cohesin. Science, 366, 1338–1345.

68. Lee, J., Chen, L.-F., Gaudin, S., Gupta, K., Spakowitz, A. and Boettiger, A.N. (2025) Kinetic organization of the genome revealed by ultra-resolution, multiscale live imaging. 10.1101/2025.03.27.645817.

69. Padovan-Merhar, O., Nair, G.P., Biaesch, A.G., Mayer, A., Scarfone, S., Foley, S.W., Wu, A.R., Churchman, L.S., Singh, A. and Raj, A. (2015) Single Mammalian Cells Compensate for Differences in Cellular Volume and DNA Copy Number through Independent Global Transcriptional Mechanisms. Molecular Cell, 58, 339–352.

70. Hildebrand, E.M., Polovnikov, K., Dekker, B., Liu, Y., Lafontaine, D.L., Fox, A.N., Li, Y., Venev, S.V., Mirny, L.A. and Dekker, J. (2024) Mitotic chromosomes are self-entangled and disentangle through a topoisomerase-II-dependent two-stage exit from mitosis. Molecular Cell, 84, 1422–1441.e14.

