## Supplementary Figures S1 to S6 for "Cell Cycle-Dependent Chromatin Motion: A Role for DNA Content Doubling Over Cohesion"

**This supplement contains:**

**Supplementary Figures S1 to S6**

**Supplementary Figure S1:**

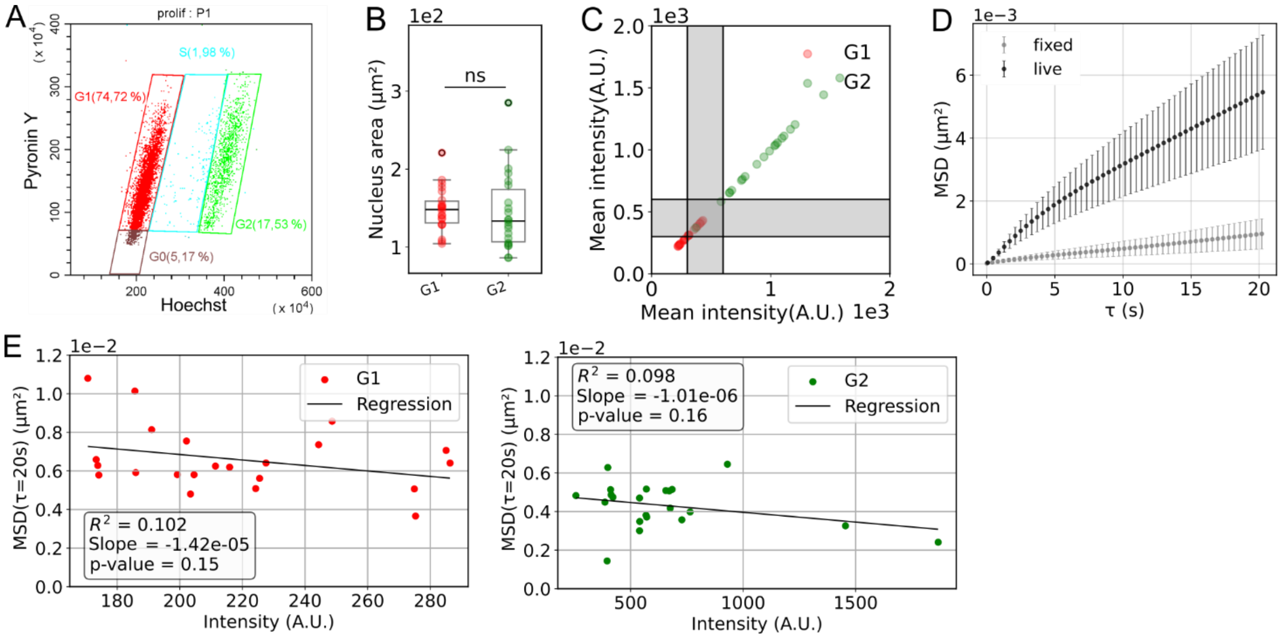


Figure S1:

A) Cell cycle distribution of an asynchronised IMR90 cell population by Fluorescence-Activated Cell Sorter obtained by quantification of DNA content (Hoechst 33324) and RNA content (Pyronin Y). The percentage of cells in each phase is indicated on the graph.

B) Nucleus area of the first frame of the analysed nuclei. Box plot description similar to Figure 1D.

C) Plot of the mean intensity. Grey bands are intensity thresholds. In this population, cells with a mean intensity smaller than 300 are exclusively in G1 phase and cells with a mean intensity larger than 600 are exclusively in G2 phase.

D) Mean ± SD of the mean MSD of live cells (N=44, same cells as in Figure 1) and para-formaldehyde fixed cells (N= 45).

E) Plots for cells in G1 (left) and G2 (right) of the mean MSD in function of the mean intensity and the respective linear regressions with p-value from Pearson correlation.

**Supplementary Figure S2:**

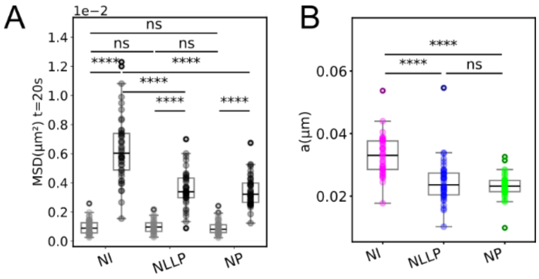


Figure S2:

A) Box plots of the mean $MSD(\tau = 20s)$ in live (black) and fixed (gray) segmented nuclei (NI, NLLP and NP). Box plot description similar to Figure 1D.

B) Box plots of the physical parameter $a$ extracted from the MSDs shown in Figure 2B in the NI, NLLP and NP. Box plot description similar to Figure 1D.

**Supplementary Figure S3:**

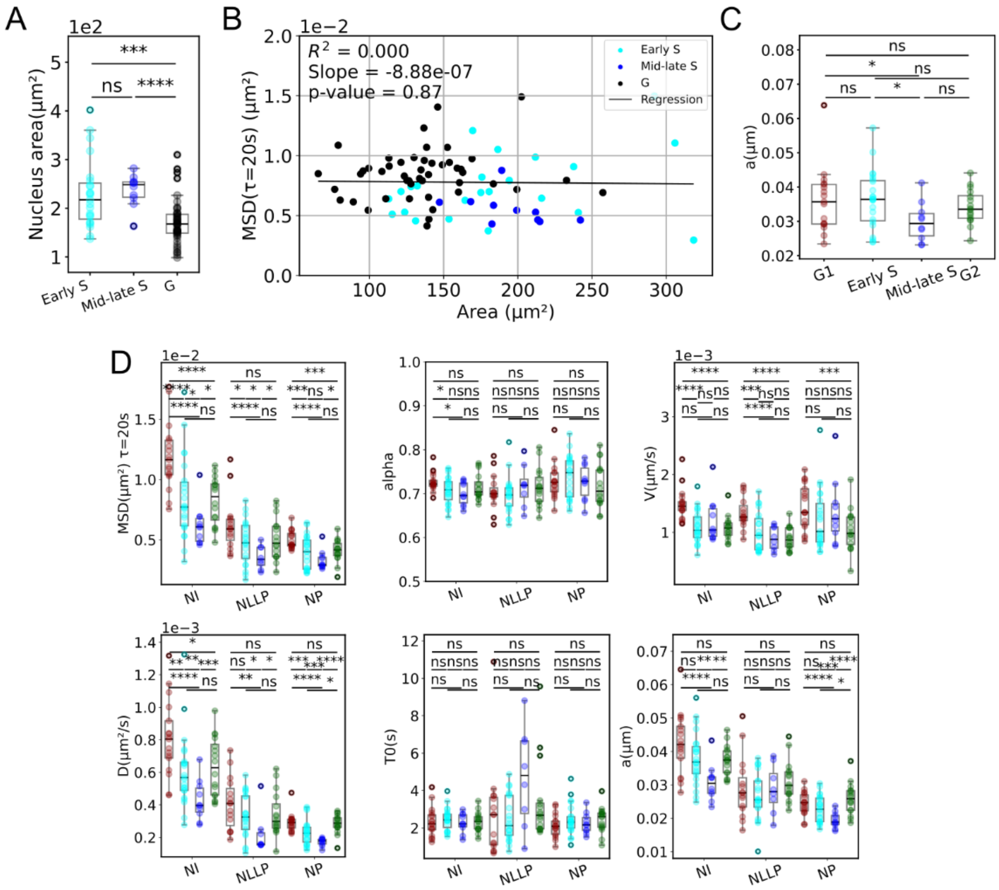


Figure S3:

A) Nucleus area of the first frame of the analysed nuclei. Box plot description similar to Figure 1E.

B) Plots for cells in G, early S and mid/late S phases of the mean MSD over the mean area and the respective linear regressions and p-value from Pearson correlation.

C) Box plots of the physical parameter (a) extracted from the MSDs shown in Figure 3D. Box plot description similar to Figure 1D.

D) Box plots of the mean $MSD(\tau = 20s)$ and the physical parameters ($\alpha, v, D, T_0$ and $a$) of G1, early S, mid/late S and G2 nuclei segmented in NI, NLLP and NP. Box plot description similar to Figure 1D.

**Supplementary Figure S4:**

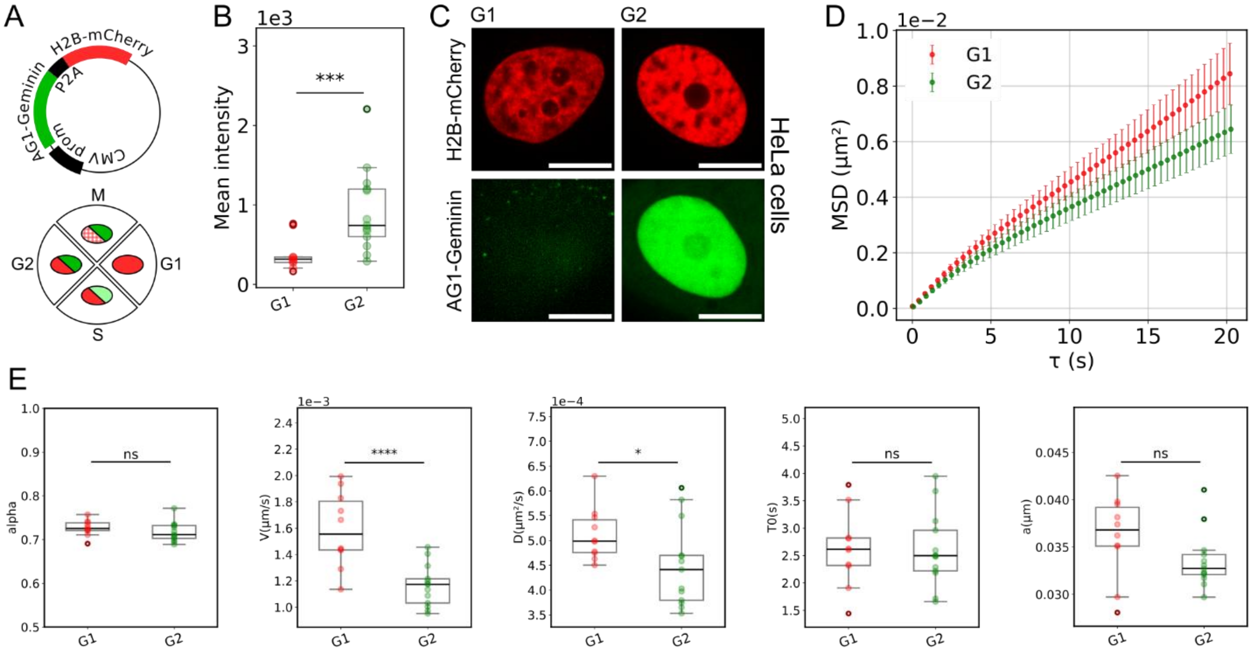


Figure S4:

A) Scheme of the FUCCI plasmid used and the consequent cell cycle distribution of the fluorescent markers.

B) Mean intensity of the first frame of the analysed nuclei. Box plot description similar to Figure 1D.

C) Representative images of G1 and G2 HeLa nuclei. Scale-bar represents 10 μm.

D) For cells in G1 (N=10, red) and G2 (N=13, green): mean±SD of the mean MSD of each nucleus.

F) Box plots of the physical parameters ($\alpha, v, D$, $T_0$ and $a$) extracted from the MSDs shown in Figure Supp 4D for cells in G1 or G2. Box plot description similar to Figure 1D.

**Supplementary Figure S5:**

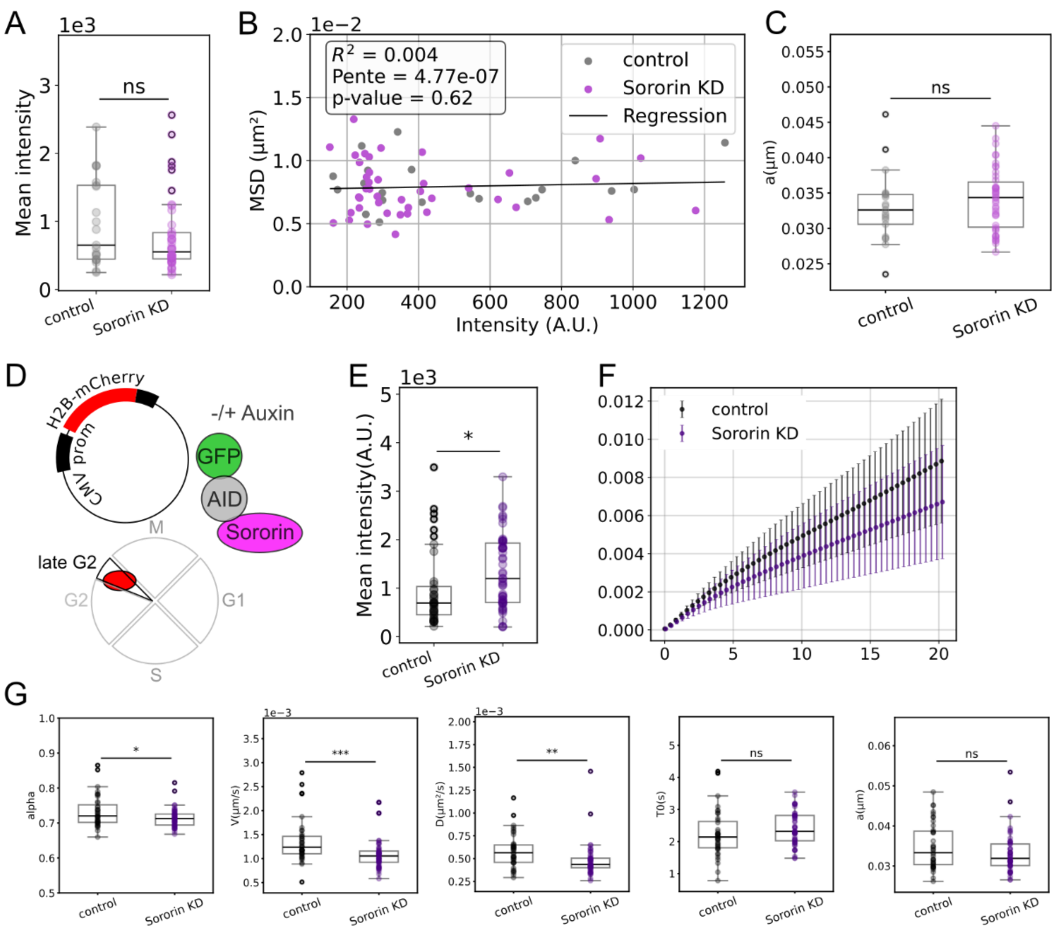


Figure S5:

A) Mean intensity of the first frame of the analysed HeLa-sororin-AID nuclei synchronised in early G2. Box plot description similar to Figure 1D

B) Plots of the mean MSD of HeLa-sororin-AID nuclei synchronised in early G2 in function of the mean intensity and the respective linear regressions with p-value from Pearson correlation.

C) Box plots of the physical parameter $a$ extracted from the MSDs shown in Figure 5C for cells synchronised in early G2 and KD or not for sororin. Box plot description similar to Figure 1D.

D) Scheme of the H2B plasmids used, the cell cycle phase studied and the cellular system used (HeLa Kyoto N-terminally-tagged sororin-AID) synchronised in late G2.

E) Mean intensity of the first frame of the analysed HeLa-sororin-AID nuclei synchronised in late G2. Box plot description similar to Figure 1D

F) For cells non-treated/control (N=39) and treated with auxin/sororin KD (N=52) synchronised in late G2: mean±SD of the mean MSD of each nucleus.

G) Box plots of the physical parameters ($\alpha, v, D$, $T_0$ and $a$) extracted from the MSDs shown in Figure Supp 5F for cells in control and sororin KD condition synchronised in late G2. Box plot description similar to Figure 1D.

**Supplementary Figure S6**

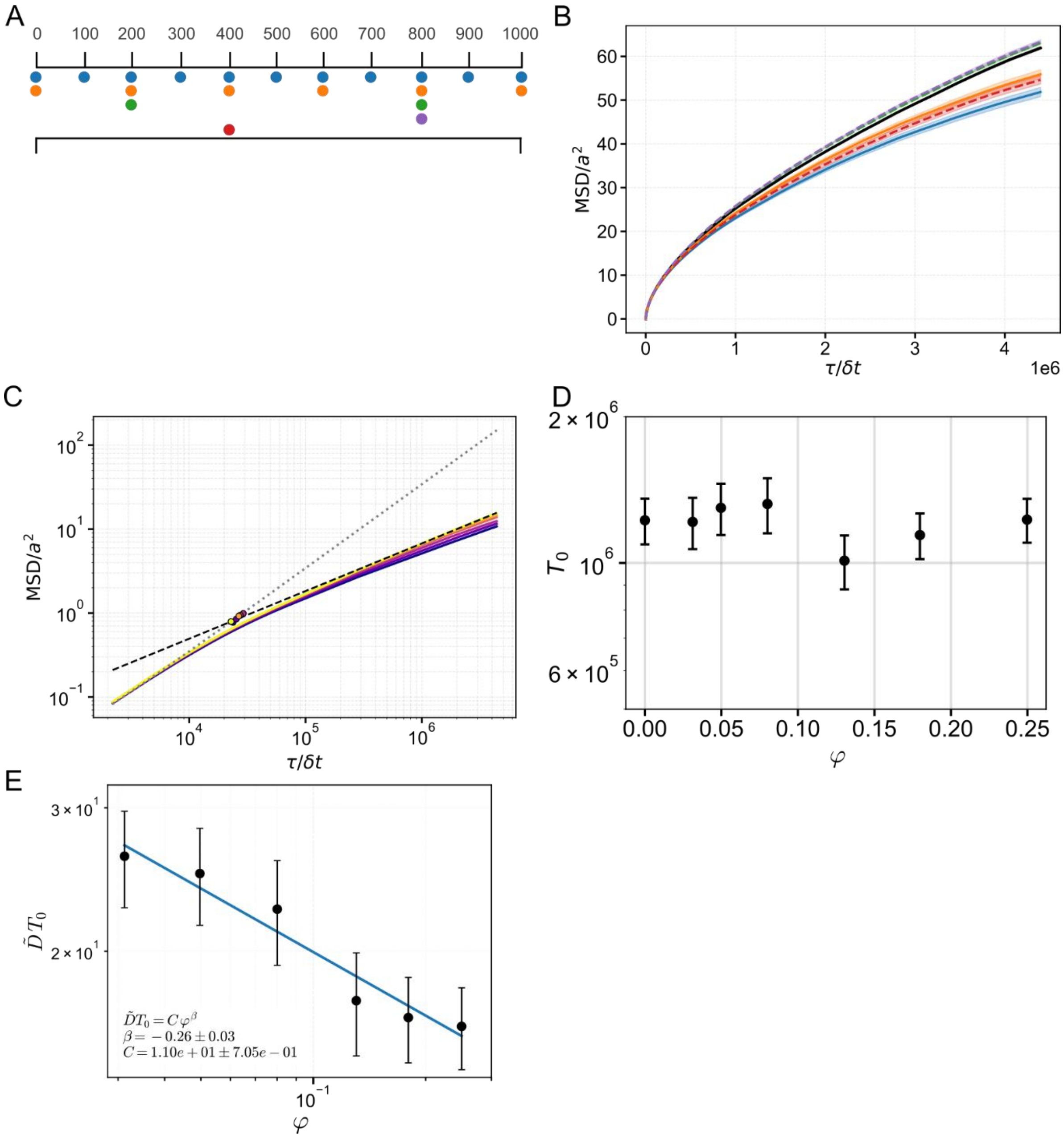


Figure S6:

A) Schematic representation of the different cohesin attachment configurations along two polymer chains of length N = 1000. The upper axis indicates the monomer index along the chains. The two black rods represent the two polymer chains. The coloured dots represent sites of attachment, i.e. the cohesive cohesin localisation. Blue markers denote regularly distributed cohesins with an attachment period of 100 monomers (one cohesin every 100 beads), orange markers correspond to an attachment period of 200 monomers. Green markers indicate a configuration with two cohesins bound at monomer indices 200 and 800. Red and purple markers represent single-cohesin configurations, with a cohesin bound at monomer index 400 (red) and 800 (purple), respectively.

B) MSD as a function of lag time for different cohesin attachment scenarios. The black curve corresponds to the control condition (no cohesin). The coloured curves correspond to the scenarios presented in B: solid lines for regularly distributed attachments and dashed ones to the other cases.
C) MSD for different volume fractions $\varphi$, ranging from $\varphi = 0.03$ (yellow) to $\varphi = 0.25$ (purple) (Figure 5F). Solid curves correspond to ensemble-averaged MSD ± SEM, dashed grey lines show linear fit $MSD(\tau) = D\tau$ at short time-scale and black dashed lines corresponds to non-linear fit $MSD(\tau) = A_\alpha \tau^\alpha$ at longer time-scale. The characteristic crossover time $\tau_0$ is estimated from the intersection of these two fits. We obtain values in the range $\tau_0 \sim (2-3) \times 10^4 \delta t$ time-steps.
D) Characteristic time $T_0$ extracted from MSD fits as a function of the volume fraction $\varphi$. The parameter $T_0$ is obtained from the fit of the MSD using the functional form $MSD = A_\alpha \tau^\alpha$.
E) Dependence of $\widetilde{DT_0}$ on the volume fraction $\varphi$. Black symbols show the values extracted from the fits of $A_\alpha$ for each confinement radius, with error bars corresponding to propagated uncertainties. The blue line shows the power-law fit $\widetilde{DT_0} = \left(\widetilde{DT_0}\right)_0 (\varphi/\varphi_0)^\beta$ with exponent $\beta = -0.26 \pm 0.03$.
